# PhISCO: a simple method to infer phenotypes from protein sequences

**DOI:** 10.1101/2022.10.23.511734

**Authors:** Ayelén S. Hernandez-Berthet, Ariel A. Aptekmann, Jesús Tejero, Ignacio E. Sánchez, Martín E. Noguera, Ernesto A. Roman

## Abstract

Although protein sequences encode the information for folding and function, understanding their link is not an easy task. Unluckily, the prediction of how specific amino acids contribute to these features is still considerably impaired. Here, we developed PhISCO, Phenotype Inference from Sequence COmparisons, a simple algorithm that finds positions associated with any quantitative phenotype and predicts their values. From a few hundred sequences from four different protein families, we performed multiple sequence alignments and calculated per-position pairwise differences for both the sequence and the observed phenotypes. We found that from 3 to 10 positions, depending on the studied case, were enough to identify positions associated with the phenotypes and perform quantitative predictions of them. Here we show that these strong correlations can be found using individual positions while an improvement is achieved when the most correlated positions are jointly analyzed. Noteworthy, we performed phenotype predictions using a simple linear model that links per-position divergences and differences in observed phenotypes. We also show that although extremely simple, predictions are comparable to the state-of-art methodologies which, in most of the cases, are far more complex. All of the calculations are obtained at a very low information cost since the only input needed is a multiple sequence alignment of protein sequences with their associated quantitative phenotype. The diversity of the explored systems makes PhISCO a valuable tool to find sequence determinants of biological activity modulation and to predict various functional features for uncharacterized members of a protein family.

## Abbreviations

PhISCO: Phenotype Inference from Sequence COmparisons, OGT: Optimal Growth Temperature, MSA: Multiple Sequence Alignment, Adk: Adenylate kinase, HIVdb: Stanford University HIV Drug Resistance Database, RDP: resistance determining positions, BR: microbial rhodopsin, ATPlid: ATP lid domain, AMPlid: AMP lid domain, HIV-PR: HIV-1 protease, SQV: saquinavir, FQV: fosamprenavir, AUC: area under the curve, MCC: Mathews correlation coefficient, ROC: Receiver Operating Characteristic.

## Statement of Significance

We explored the association between protein sequences and quantitative phenotypes to find residues responsible for phenotype modulation. To this end, we used a simple measure based on the per-position correlation between sequence divergence and phenotype difference. Several function-linked positions were identified in four case studies corresponding to different protein families. Additionally, we used the most highly correlated positions to make quantitative predictions. This extremely simple strategy was useful to identify candidate residues for mutagenesis or other analysis in sequence-function studies and also, for the phenotype prediction of uncharacterized proteins or organisms.

### Introduction

Proteins are essential molecules for cellular metabolism. Conformational dynamics, stability and functional features such as catalytic efficiency or interaction capability are encoded in their sequences. Thus, this information must be accessible from analysis of protein sequences. As an example, coevolutionary analysis from sequences has succeeded in the prediction of native contacts and their concomitant structures with the aid of statistical and bioinformatic tools [1]. Although structure and function are tightly linked, the prediction of functional features in high detail is still considerably impaired.

There is great interest in understanding the mechanisms through which homologous proteins are adapted to the very heterogeneous selective pressures to which they are subjected. A deep analysis of these mechanisms could provide pathways to modulate phenotypic characteristics such as stability, chromophore absorption, catalytic efficiency and ligand affinities. From an applied point of view, this knowledge can be used to design proteins whose properties are optimized to work in the harsh conditions frequently found in industrial processes such as high temperatures or increased salt concentrations. Whereas genetic variation allows the phenotypic variation, a delicate balance of molecular interactions is necessary to preserve the proper stable three-dimensional structure in a functional competent state and consequently, not all the residues across the protein sequence are equally available to functional tuning.

In this sense, variations in the sequences may be reflected in variations in observed phenotypes. However, usually, the changes in the observed phenotypes do not appear in an additive way (’additive’ means that the effect of each individual mutation accumulates as a simple sum of effects). In many cases, accumulation of mutations does not have a linear counterpart in the observed trait. This type of phenomena is called epistasis and usually named as a non-additive effect of mutations. A reasonable hypothesis is mentioned in Otwinowski J. et al. posing that mutations have an additive effect in an unobserved trait which is related in a non-linear way to the observed trait [9].

Multiple existing algorithms aim at the quantitative prediction of phenotypes. Some of them take a protein structure as a starting point, while others are sequence-based. Structure-based predictions of protein stability are time-consuming, require expert supervision, and are moderately accurate [2]. Although with the aid of structure predictors as AlphaFold2 the structure-requirement is no longer an obstacle for most targets, predictions of functional properties are still not always performed with significant accuracy. Sequence-based prediction algorithms attempt to associate amino acid abundances and/or physical properties at specific sequence positions with phenotype values such as the optimal working temperature of a protein/organism [3-5], the impact of a mutation in disease [6], binding affinity [7] or the spectroscopic properties of a protein [8]. Since the nature of the problems tackled is quite diverse, it comes to no surprise that individual methods are tailored towards specific questions. As a consequence their applicability is very limited [9]-[10]. Their accuracy is quite diverse ranging from moderately low to high and many of these methods are based on machine learning techniques requiring extensive training data sets and suffer from parameter interpretation problems [11].

In this work we propose a simple method that we called PhISCO from Phenotype Inference from Sequence COmparisons to 1) identify protein positions correlated with the acquisition of specific phenotypes and, from these findings, 2) quantitatively predict phenotypes for a target protein sequence. The idea behind this method is that only a limited set of residues is responsible for modulation of functional phenotypes in a protein family, it is expected that protein variants working in similar conditions exhibit similar residues at these positions. To search these residues, we performed calculations to correlate per-position sequence differences (“sequence divergence”) and the corresponding phenotype differences of the involved sequences. At this point, two considerations are important to support this notion: i) the phylogenetically distant organisms would obtain similar residues due to a convergence phenomenon and ii) these positions would have a “dominant” effect acting in an independent manner from the sequence background (epistasis). Given the absence of epistasis, the historical contingency in phenotype modulation would be absent. Then, although different lineages could develop independent adaptation strategies, the same sequence changes would alter the phenotype in a similar way regardless of the lineage in which they operate.

We illustrate the performance of our method using four case studies of protein families with different associated properties: a set of adenylate kinases from Archaea with their associated OGT which has been studied by several methods, a set of microbial rhodopsins whose retinal cofactor absorption occurs at different wavelengths [8], a set of myoglobin sequences from mammals and their maximal concentration in muscle, and the potency of HIV protease inhibitors for clinical isolates [13]. To our knowledge, only a few methods exploit the sequence information of different organisms with a simple metric that relates the sequence similarity to the proximity of the OGT [12]. Our proposal minimizes calculation requirements with almost no assumption on a minimal set of parameters.

### Results

## PhISCO overview

In this work we developed an algorithm that calculates per-position sequence divergences in the alignment as well as the associated phenotype differences (Figure 1A). Divergences are calculated using an identity matrix, where values of 0 or 1 are assigned to identical or different amino acids, and phenotype differences are treated as the corresponding absolute values (ΔPhenotypes in Figure 1 and equations thereof). Association between these two magnitudes is evaluated position by position using the Spearman rank-order correlation coefficient (ρ) [37], from which the positions more correlated with phenotypes can be extracted and studied. In addition, a set of positions highly correlated with the phenotypes can be used for the prediction of the phenotype value of uncharacterized sequences. For this end, we developed a specific heuristic (Figure 1B) to explore the existence of a set of positions that, when analyzed jointly, renders a higher correlation than when individually analyzed. Briefly, the correlations between sequence divergence and phenotype differences are calculated for sets of positions that increase in size by the addition of one position at a time, starting from the most correlated individual position up to the least. Thus, at the beginning of the calculation, the ρ- value corresponds to the position with highest correlation; in the next step, it sums the divergence of the second highest correlated position to that of the former position and recalculates the ρ-value. The process is repeated until all of the positions are included (Figure 1B).

**Figure 1.**
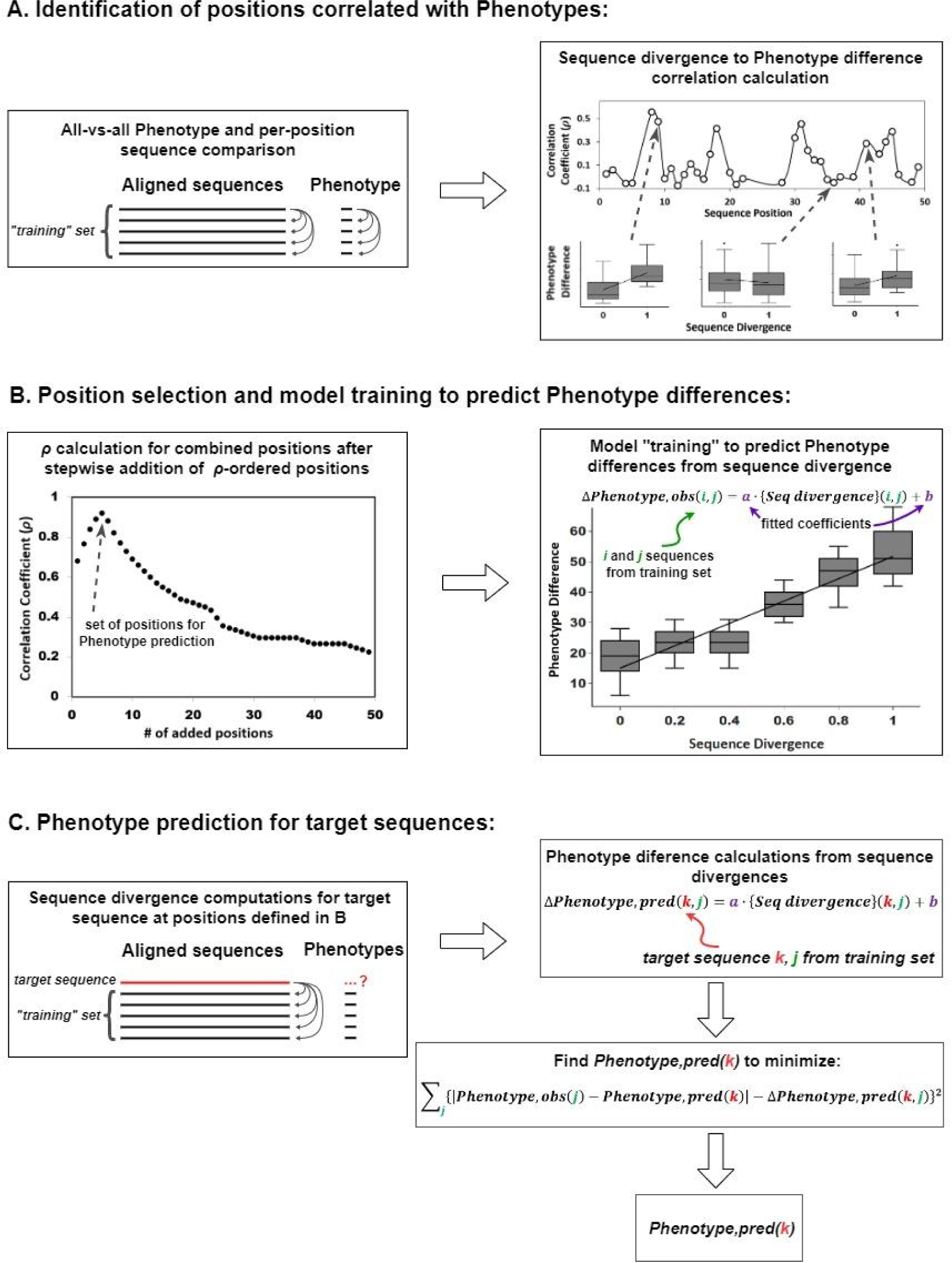
Detailed workflow of PhISCO algorithm. A. Identification of positions correlated with the phenotypes. Sequences from a protein family were aligned and pairwise similarity comparison was performed for the sequence and the observed features, for each position. A Spearman’s rank-order correlation analysis was performed and rho values were plotted for each position. B. Position selection and model training to predict phenotype differences. Positions were ordered from highest to lowest *ρ*- values and the correlation was recalculated by sequentially adding the divergence of each position. Thus, at the beginning of the calculation, the *ρ*-value corresponds to the position with highest correlation and the last point corresponds to the whole sequence. C. To do the prediction the target feature was chosen as the one that minimizes the squared-sum of the difference between the predicted ΔPhenotype from the linear model and the difference between the unknown and each phenotype in the training set sequences. The calculation was repeated for all of the sequences of the validation set.

The group of positions with highest ρ-value are then used for phenotype prediction of target sequences. For this end the training sequences (those with already known phenotypes) are used for pairwise comparisons as described above using the group of positions with highest ρ-value, and the correlation between sequence divergence and ΔPhenotype is calculated using a linear equation (Figure 1C). The equation coefficients derived from the linear regression are then used to extrapolate the ΔPhenotype from the divergence values at the specified positions of the target sequence to those of the training set. Finally, the target phenotype is derived from minimization of a sum of squares calculated from extrapolated ΔPhenotypes and those calculated from the difference in the unknown (target) phenotype and the known phenotypes of the training set. A detailed description of the minimization procedure and equations is included in the Methods section. In the rest of the results section we discuss the application of this algorithm to four studied cases and analyze the results.

## The case of archaeal adenylate kinases (*ADK*)

Enzymes involved in key-metabolism steps need to properly work in conditions similar to those in which the organisms grow [14]. One of these key enzymes is AdK, which plays a central role in energy metabolism by regulating the ATP concentration [14,15]. We focused on AdK from *Archaea*, since these organisms grow in a wide range of temperatures and there is abundant information about sequences and growing conditions. To constrain the analysis to sequences from organisms subjected to similar adaptive strategies we performed a clusterization analysis using CLANS as described in Methods section. We found two groups of sequences: one of 163 and another of 80 sequences (**Figure 2A**). While the minor cluster contains proteins that had been reported as trimeric [16–18], none of the sequences from the major cluster has a reported structure. However, the high identity with bacterial monomeric homologs suggests that this set may also be monomers. In that sense, and since these proteins seem to have different oligomeric states that can redound in differences in their catalytic mechanisms, we focus on the major cluster. An analysis of the whole sequence divergence and their associated *ΔOGT* correlation shows a *R^2^* = 0.623 ± 0.008 (**Figure S1**). However, a region is evident where although divergence is high, the *ΔOGT* is low. To explore whether there are specific positions that exhibit a tighter link to *ΔOGT* than the whole sequence, we studied the per-position correlation as described in *Methods*. Spearman’s rank correlation coefficients showed a widespread distribution with a mean value of 0.17 ± 0.21 (**Figure 2B**). We evaluated if there exists a group among these positions that further improve the correlation with *ΔOGTs* by correlation recalculation after sequential addition of *ρ*-value ordered positions. Whereas this method to explore the joint correlation is not exhaustive, it provides a path with high chance to find a set of residues that captures most of the sequence-phenotype association. For the case of AdK, we found three positions that combined yielded the highest *ρ*-value (0.83) (**Figure 2C-E**, **Table 1**). We mapped these positions in the AdK X-ray structure (PDB ID 1AKE) (**Figure 2D**). All of them are located at the ligand binding sites: while position 54 (38 in *E.coli* sequence, Table 1) is located in the AMP lid domain (AMPlid), positions 160 (116 in *E.coli* sequence, Table 1) and 221 (175 in *E.coli* sequence, Table 1) are located in the ATP lid domain (ATPlid) [19]. Computational analysis of coevolution combined with molecular dynamics simulations showed that residue 38 is highly coupled to spatially distant positions [20]-[21]. Regarding positions in the ATPlid, when the mutation of residue 116 from I to G is introduced in the *E. coli* AdK ATP binding is improved without significantly protein stability effect but decreasing catalytic efficiency [19]. Also, structure-based simulations and normal mode analysis showed that both residues 116 and 175 are involved in local unfolding events with effects in AdK catalysis [21,22]-[23]. Given that this reduced set of positions are involved in enzyme functionality and captures most of the correlation, we next tested its capability to predict OGT. The sequence dataset was split in training and validation sets (50 % of randomly picked sequences each one). This random sampling was repeated 50 times to evaluate prediction robustness. A representative prediction plot with the mean *R*^2^ value is shown in **Figure 2F**. Interestingly, the prediction using the three positions gave more accurate results (*R^2^ = 0.85*) than the whole sequence (*R^2^ = 0.56*). This finding, besides showing that those positions are linked to OGT, makes this algorithm very helpful for OGT prediction based on AdK sequences.

**Figure 2.**
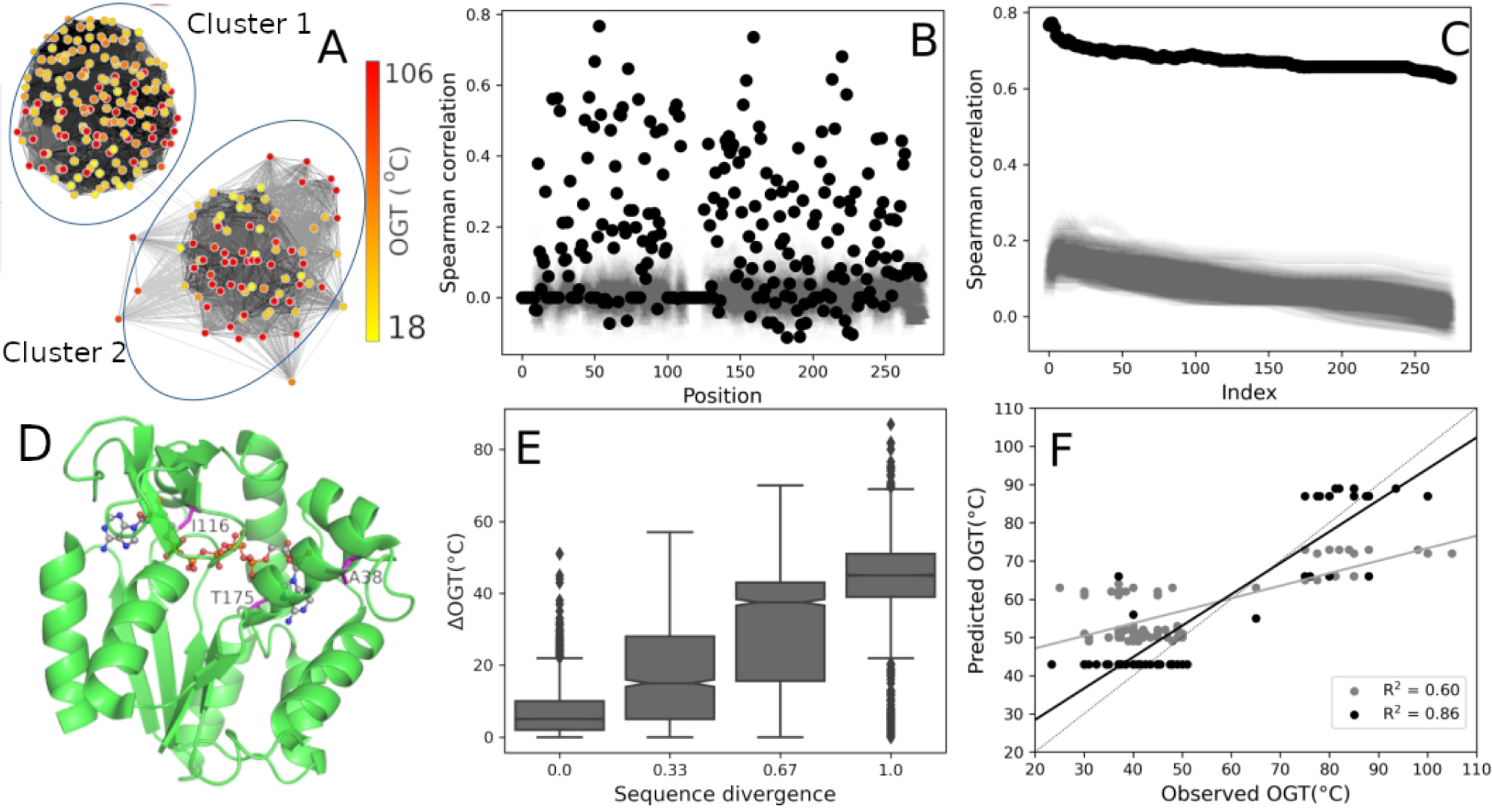
Optimal Growth Temperatures associated with Archaeal Adenylate Kinase (AdK). A. Sequence clusterization according to pairwise similarity threshold. Cluster 1 has 163 sequences while cluster 2 has 80. B. Spearman rank-ordered correlation coefficients per position in the MSA (black filled circles) and control calculations after randomization of phenotype values (100 rounds of random assignments of phenotypes to sequences, grey points). C. Joint correlation coefficients of the AdK by addition of positions ranked in a descendent order (black filled circles) or by adding the same positions in the same order but after their phenotype values have been randomly permuted (100 rounds, gray points). D. Positions found in the joint correlation analysis mapped in a high resolution structure (PDB ID 1AKE). E. Distribution of the phenotype differences at different sequence divergence values. (R2 = 0.86 from a linear fit of ΔOGT vs sequence divergence). F. Prediction of OGT from sequences for the most correlated positions (black) and the entire sequence (gray).

**Table 1.**
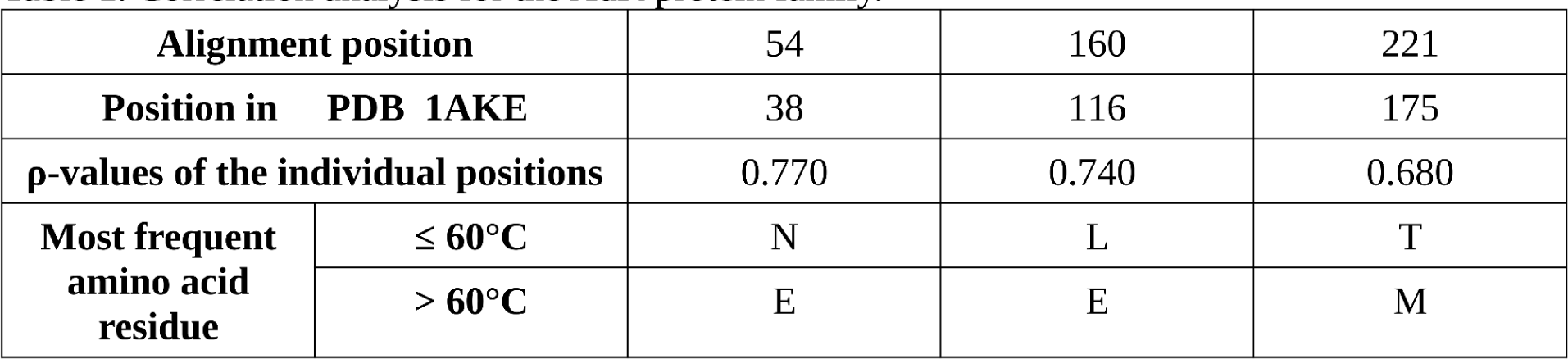
Correlation analysis for the ADK protein family.

## The case of microbial rhodopsins (BR)

To study if PhISCO can be used in other scenarios, we performed calculations on a diverse set of microbial rhodopsins. They belong to a ubiquitous photoreceptive membrane protein family that has diverse functions as light-driven ion pumps, light-gated ion channels, photochromatic gene regulators and light-regulated enzymes. These proteins bind all-trans retinal molecules via a protonated Schiff-base linkage and exhibit a variety of specific visible absorption wavelengths. This wide-range color tuning of the retinal is considered to be achieved by optimizing the steric and/or electrostatic interaction with surrounding amino-acid residues [8]. Based on a previous work by *Karasuma et al*.[8] and *Inoue et al*. [24] we evaluated whether our method was able to identify positions associated with the fine tuning of the absorption-wavelength of microbial rhodopsins. To this end, we worked with the published data containing microbial rhodopsin sequences and associated wavelengths [8]. Analysis of the sequences using CLANS showed three main clusters with similar numbers of elements (**Figure 3A**). The least populated group showed an extremely narrow distribution of absorption wavelengths while the other two spanned broader ranges. Analysis of the major cluster revealed sequences ∼30- 60% identical that span a distribution of *ΔWavelengths* from 25 nm to 125 nm with a correlation of *R^2^* = 0.279 ± 0.004 (**Figure S1**). To seek positions that are more tightly linked to the phenotype, we studied the per-position *ρ*-value. The maximum value was 0.47, while the analysis of the joint correlation shows a peak when we used the ten most correlated positions (*ρ*-value = 0.65) (**Figure 3C-E and Table 2**). We mapped these positions in the crystal structure of *Halobacterium salinarum* (PDB ID 1C3W). Within them, positions 71 (85 in *H. salinarum* sequence, Table 2) and 104 (118 in *H. salinarum* sequence, Table 2) are in direct contact to the retinal molecule [8]. Modification of some loops in the Br from *H. salinarum*, involving positions 87, 58 and 51 (101, 68 and 61 in *H. salinarum* sequence, respectively, Table 2), have been reported to cause a maximal absorption wavelength shift [25]. The remaining positions are not in direct contact to retinal, however, positions 3, 10 and 12 (10, 19 and 21 in *H. salinarum* sequence, respectively, Table 2) are in direct interaction with the lipids located in the inter-monomer space. Noteworthy, mutation of position 10 was reported to have effects in maximal-absorption wavelength [26]. The sequence dataset was split in training and validation sets (50 % of randomly picked sequences each one). This random sampling was repeated 50 times to evaluate prediction robustness. A representative prediction plot with the mean R^2^ value is shown in **Figure 3F**. The chosen set of positions improved the correlation (*R^2^ = 0.72*) with respect to the whole sequence (*R^2^ = 0.60*) suggesting that a small subset of positions is rich in phenotypic information and can be easily extracted using PhISCO.

**Figure 3.**
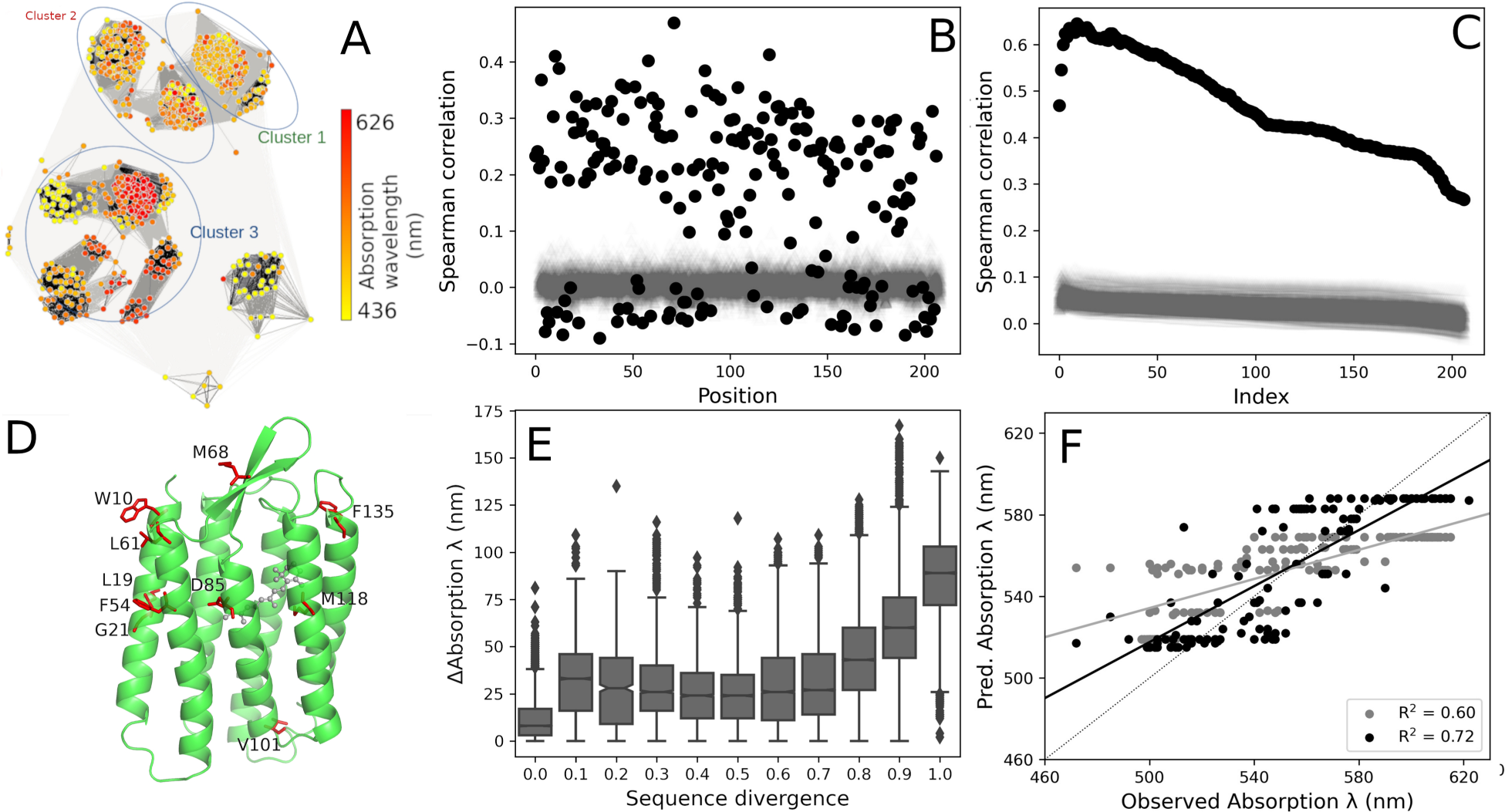
Maximum absorption wavelength of variants of bacterial rhodopsins (Br). A. Sequence clusterization according to pairwise similarity threshold. Cluster 1 has 284 sequences, cluster 2 has 281 sequences and cluster 3 has 245 sequences. B. Spearman rank-ordered correlation coefficients per position in the MSA (black filled circles) and control calculations after randomization of phenotype values (100 rounds of random assignments of phenotypes to sequences, grey points). C. Joint correlation coefficients of the microbial rhodopsins by addition of positions ranked in a descendant order (black filled circles) or by adding the same positions in the same order but after their phenotype values have been randomly permuted (100 rounds, gray points). D. Positions found in the joint correlation analysis mapped in a high resolution structure (PDB ID 1C3W). E. Distribution of the phenotype differences at different sequence divergence values (R2 = 0.58 from a linear fit of ΔAbsorption wavelength vs sequence divergence). F. Prediction of absorption wavelength from sequences for the most correlated positions (black) and the entire sequence (gray).

**Table 2.**
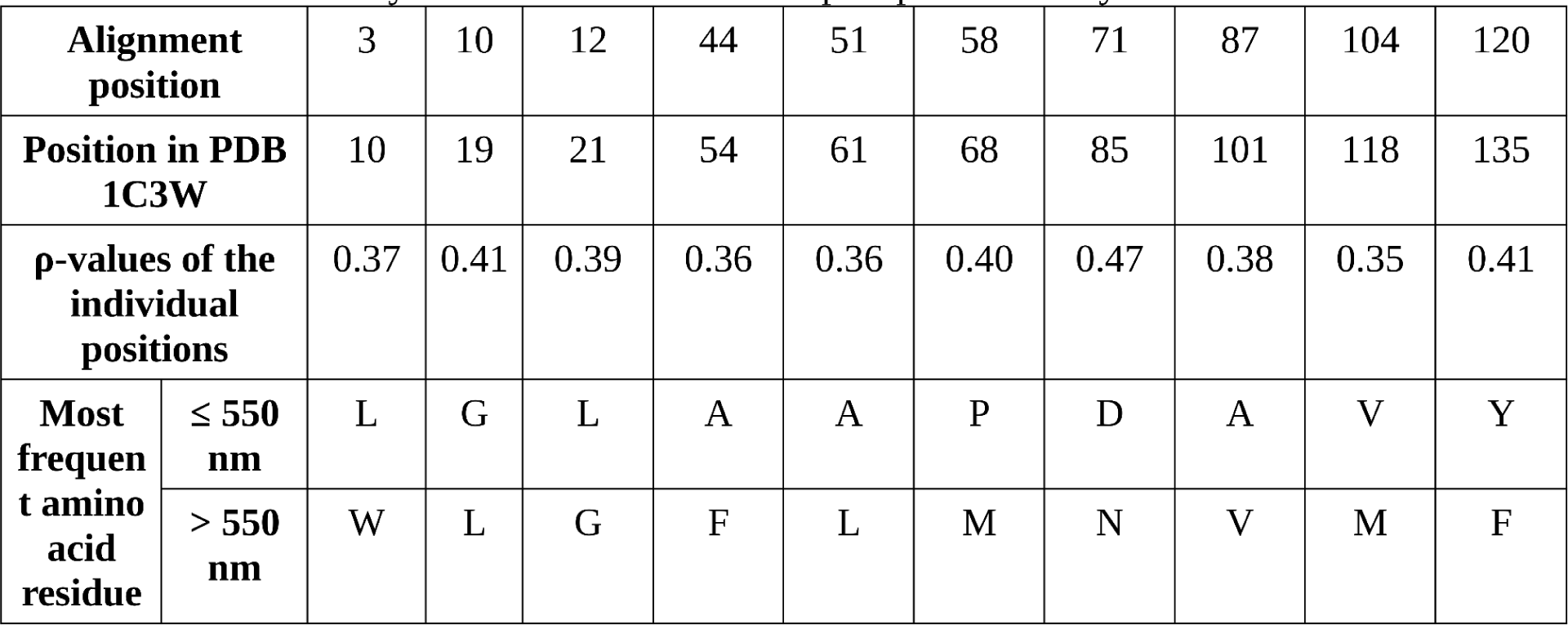
Correlation analysis for the microbial rhodopsin protein family.

## The case of mammalian myoglobin and its maximal muscle concentration

Aquatic mammals such as whales and seals have evolved mechanisms to hold their breath for periods of over an hour [27]-[28]. Mammals’ diving capacity is tightly linked to the myoglobin concentration in muscle, which can be over 30-fold higher in mammalian elite divers as compared to land species [28,29]-[30]. In order to increase protein concentration, mammalian myoglobins have undergone extensive modifications to enhance solubility without triggering aggregation processes. Strategies to increase solubility highly rely on the increase of the net surface change [29], although alternative mechanisms such as increasing thermodynamic stability and hydrophobic patch shielding are also involved [31]. To evaluate positions that are linked to changes in myoglobin concentration, we studied a set of 65 mammalian myoglobin sequences from terrestrial and aquatic mammals with available myoglobin concentration data [29,31]. An inspection of the sequences using CLANS showed a single cluster (**Figure 4A**). The pairwise identity of these sequences are between 80% and 90% spanning a range of two orders of magnitude in the Δ*log10[Mb]* with no obvious correlation between them (*R^2^* = 0.241 ± 0.001) (Figure S1). To evaluate if there are positions linked to the protein concentrations in the different organisms we studied per-position correlation. Most of the positions were invariant among sequences, then there are only few positions that had higher than zero *ρ-value* (mean value of 0.03 ± 0.07) (**Figure 4B**). The highest individual *ρ-value* was 0.41, while the most correlated positions (62, 101 and 152), when jointly analyzed, improved the *ρ-value* to 0.51 (**Figure 4C-E**). The mutation Q152H is almost exclusively found in diving mammals [29]. A correlation between thermodynamic stability and myoglobin cell concentration has been reported [31]-[32]. Studies of the evolution of myoglobin from terrestrial mammals to the elephant seal note mutations in 18 positions. Among them, PhISCO identified positions 62, 101 and 152 [31]. Notably, 152 and 101 side chains are close in the reported structures, such as PDB ID 1ABS (Figure 4D) (Table 3). To perform *log10[Mb]* prediction, the sequence dataset was split in training and validation sets (50 % of randomly picked sequences each one). This random sampling was repeated 50 times to evaluate prediction robustness. A representative prediction plot with the mean R^2^ value is shown in **Figure 4F**. Prediction of the *log10[Mb]* using these positions yielded a *R^2^ = 0.61* while using the whole sequence it was *R^2^* = *0.44*.

**Figure 4.**
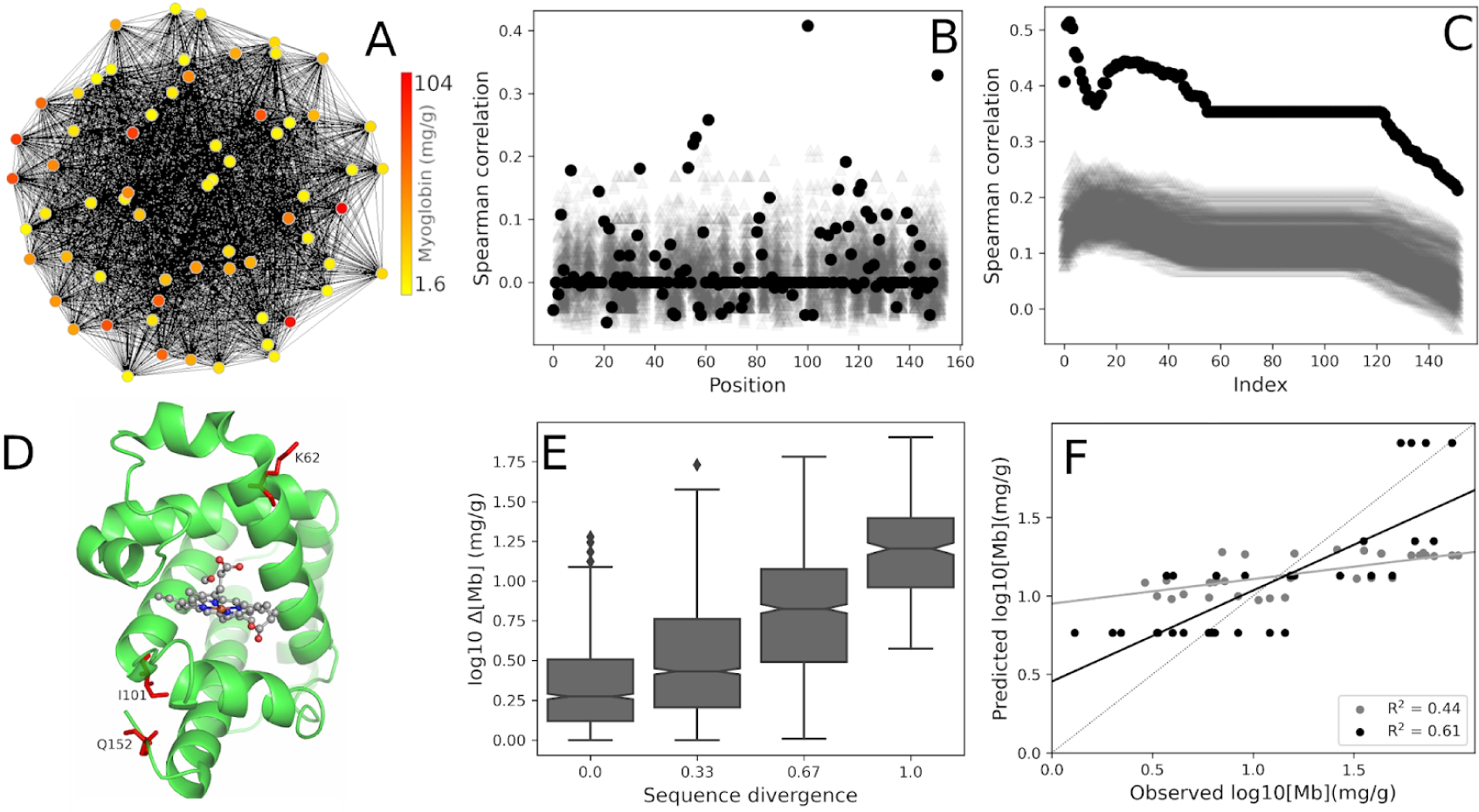
Myoglobin muscle concentrations in mammals (mmMb). A. Sequence clusterization according to pairwise similarity threshold. B. Spearman rank-ordered correlation coefficients per position in the MSA (black filled circles) and control calculations after randomization of phenotype values (100 rounds of random assignments of phenotypes to sequences, grey points). C. Joint correlation coefficients of the mmMb by addition of positions ranked in a descendent order (black filled circles) or by adding the same positions in the same order but after their phenotype values have been randomly permuted (100 rounds, gray points). D. Positions found in the joint correlation analysis mapped in a high resolution structure (PDB ID 1ABS). E. Distribution of the phenotype differences at different sequence divergence values (R2 = 0.55 from a linear fit of Δ log10[Mb] vs sequence divergence). F. Prediction of mmMb from sequences for the most correlated positions (black) and the entire sequence (gray).

**Table 3.**
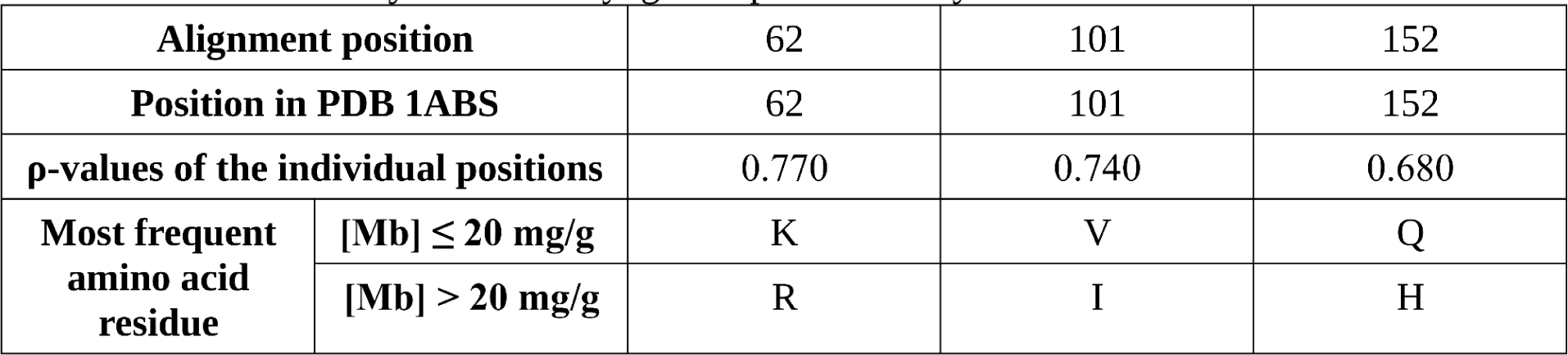
Correlation analysis for the myoglobin protein family.

## The case of HIV-protease and inhibition potency in clinical isolates

We tested the capability of our algorithm to predict sequence positions associated with antiviral drug resistance in HIV-1 strains. This phenomenon results mainly from prolonged patients-exposure to drug treatments, leading to the fixation and widespread distribution of resistant strains. We focus on the HIV-1 protease (HIV-PR), which is one of the most structurally characterized viral enzymes. HIV-PR is the target of several drugs that are analogs to the transition state of the peptide substrate. Mutations of clinical relevance are observed in around 30% of residues in the HIV-PR sequence, we call those positions resistance determining positions (RDP). We studied the case of two inhibitors: fosamprenavir (FPV) and saquinavir (SQV). Resistance to these inhibitors is associated with different sets of RDP, with some overlapping (Table 4). Using PhISCO with the sequences recovered from HIVdb with their associated inhibition potency for each drug, we found the correlation between positions and the studied phenotype. PhISCO successfully identified most of the RDP for both inhibitors. In the top 20 highest scoring positions, we found 14 out of 16 RDP for FPV and 12 out of 15 for SQV (**Supplementary Table 1**). From the set of RDP relevant for FPV and SQV (**Table 4**) [33], Some of the identified positions are in direct binding to the inhibitor, as observed in positions 47, 50, 82 and 84. Mutations in these positions are mainly I84V, I47V, I50V and V82A, where substitutions result in loss of van der Waals interactions with the inhibitor. This is also the case for position 48 where substitution from G to V causes the valine side chain to impede forming key interactions with the drug. RDP involving more indirect effects were also captured, as involved in substitutions positions 54 and 73, where structural changes are propagated to active site residues, and positions 24, 50 and 53 where structure or stability of the inter-subunit interface is compromised, affecting inhibitor binding.

**Table 4.**
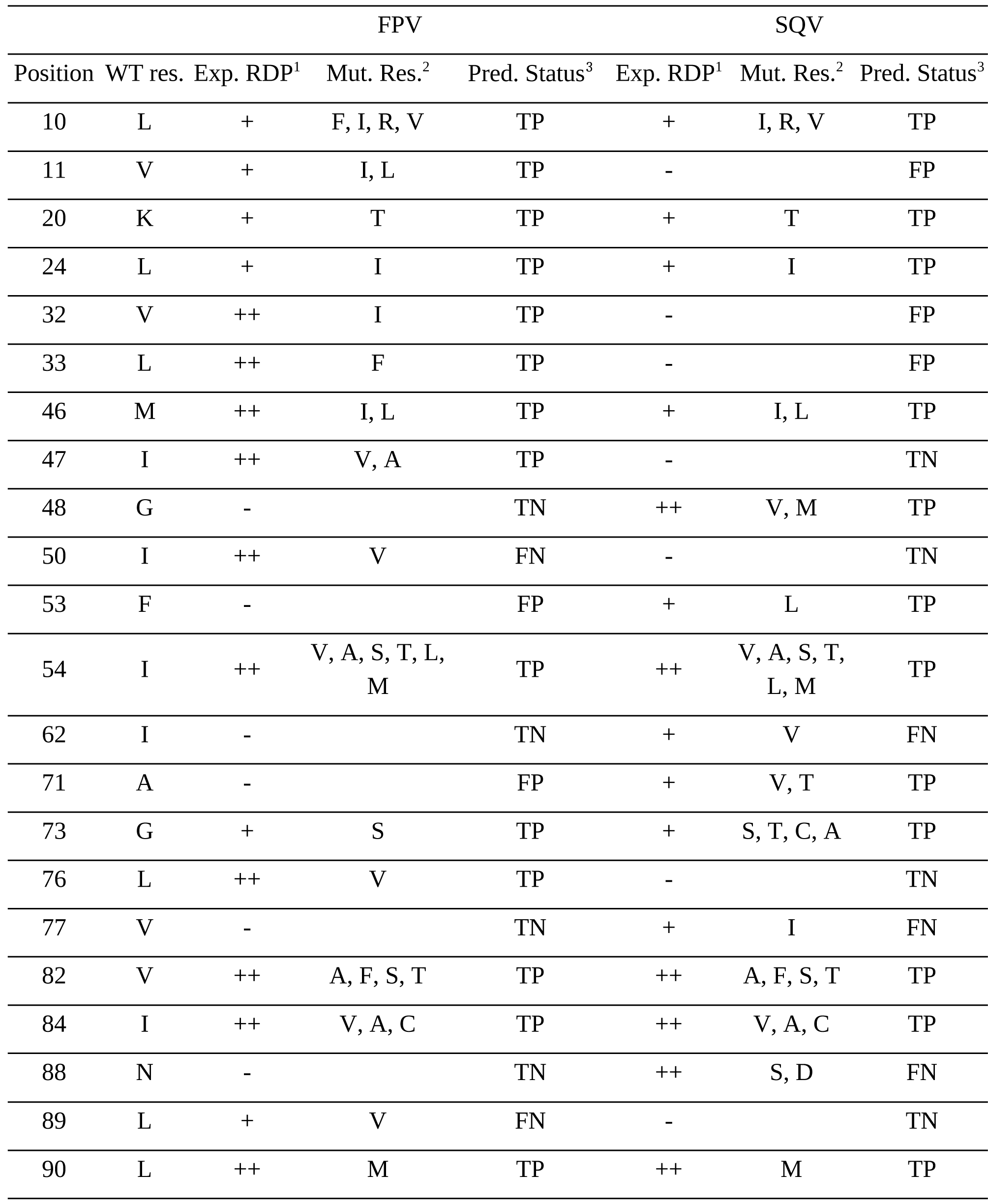
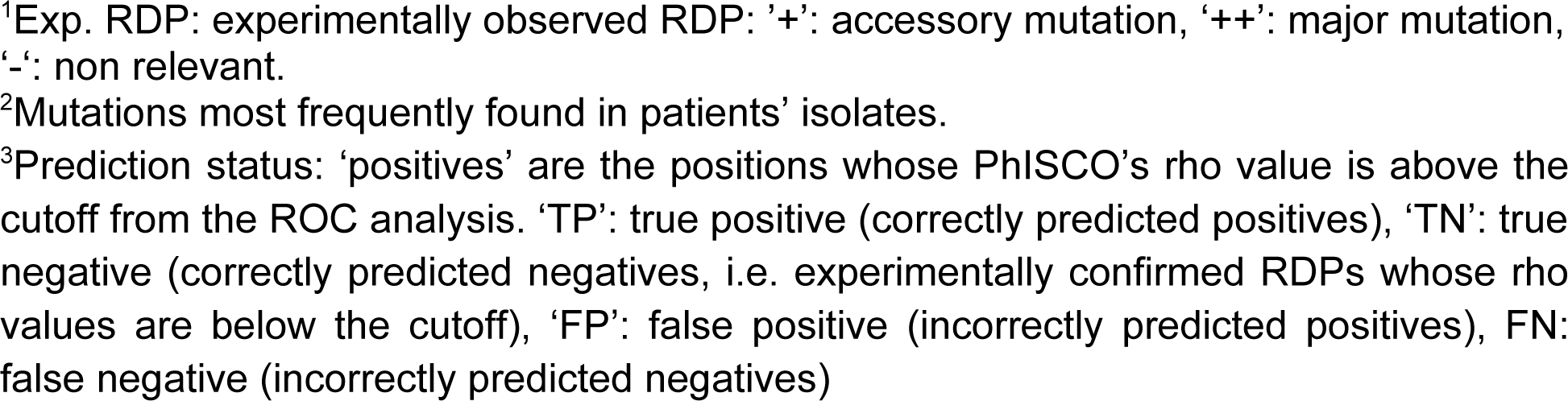
Prediction of resistance associated positions in HIV-PR.

| Position | WT res. | FPV |  |  | SQV |  |  |
| --- | --- | --- | --- | --- | --- | --- | --- |
|  |  | Exp. RDP<sup>1</sup> | Mut. Res.<sup>2</sup> | Pred. Status<sup>3</sup> | Exp. RDP<sup>1</sup> | Mut. Res.<sup>2</sup> | Pred. Status<sup>3</sup> |
| 10 | L | + | F, I, R, V | TP | + | I, R, V | TP |
| 11 | V | + | I, L | TP | - |  | FP |
| 20 | K | + | T | TP | + | T | TP |
| 24 | L | + | I | TP | + | I | TP |
| 32 | V | ++ | I | TP | - |  | FP |
| 33 | L | ++ | F | TP | - |  | FP |
| 46 | M | ++ | I, L | TP | + | I, L | TP |
| 47 | I | ++ | V, A | TP | - |  | TN |
| 48 | G | - |  | TN | ++ | V, M | TP |
| 50 | I | ++ | V | FN | - |  | TN |
| 53 | F | - |  | FP | + | L | TP |
| 54 | I | ++ | V, A, S, T, L,<br>M | TP | ++ | V, A, S, T,<br>L, M | TP |
| 62 | I | - |  | TN | + | V | FN |
| 71 | A | - |  | FP | + | V, T | TP |
| 73 | G | + | S | TP | + | S, T, C, A | TP |
| 76 | L | ++ | V | TP | - |  | TN |
| 77 | V | - |  | TN | + | I | FN |
| 82 | V | ++ | A, F, S, T | TP | ++ | A, F, S, T | TP |
| 84 | I | ++ | V, A, C | TP | ++ | V, A, C | TP |
| 88 | N | - |  | TN | ++ | S, D | FN |
| 89 | L | + | V | FN | - |  | TN |
| 90 | L | ++ | M | TP | ++ | M | TP |

<sup>1</sup>Exp. RDP: experimentally observed RDP: '+' : accessory mutation, '++': major mutation, '-': non relevant.
<sup>2</sup>Mutations most frequently found in patients' isolates.
<sup>3</sup>Prediction status: 'positives' are the positions whose PhISCO's rho value is above the cutoff from the ROC analysis. 'TP': true positive (correctly predicted positives), 'TN': true negative (correctly predicted negatives, i.e. experimentally confirmed RDPs whose rho values are below the cutoff), 'FP': false positive (incorrectly predicted positives), FN: false negative (incorrectly predicted negatives)

The availability of abundant data regarding the resistance positions for HIV-PR allowed us to examine the performance of our method using the Receiver Operating Characteristic (ROC) curve and their respective Area Under the Curve (AUC, **Figure 5**). We combined our scores with available data of RDP for each inhibitor, resulting in AUC values of 0.94 and 0.89 for FPV and SQV, respectively. From the 16 experimental RDP for FPV, 14 ranked above the cutoff value taken from the maximal Youden’s index of the ROC curve. Globally, our method attained an accuracy for RDP prediction of 91 percent, with a Mathews correlation coefficient (MCC) of 0.75. In the case of SQV, 12 of a total of 15 RDP ranked above the cutoff, with an accuracy of 90 percent and MCC 0.69. Noteworthy, for both inhibitors, PhISCO analysis of the positions joint-correlation showed a peak for the first 7 (*ρ-values* of 0.77 and 0.73 for FPV and SQV, respectively). All of them are described as RDP (**Figure S2**). To predict the levels of resistance we divided the input sequences in training and validation sets by randomly choosing 50% of the sequences for the training set and the rest for the validation set (repeated 50 times). We observed correlation between predicted and observed values of *R*^2^ = 0.60 and 0.52 for FPV and SQV, respectively. These values are higher than those obtained using the whole HIV-PR sequence, which renders *R*^2^ = 0.40 and 0.31 for FPV and SQV, respectively (**Figure S2**).

**Figure 5.**
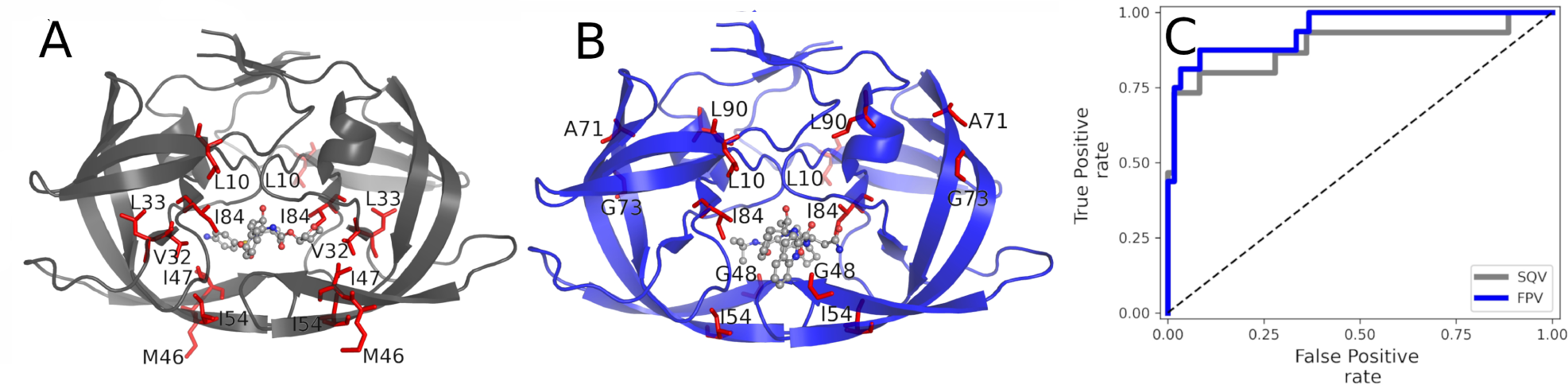
Predicted positions in HIV-PR sequence associated with inhibitor resistance. Most correlated positions (red sticks) for resistance to: A. FPV (mapped in the crystal structure for the complex with amprenavir, the active metabolite of FPV) or B. SQV (mapped in the crystal structure for the complex with this molecule). C. ROC analysis relating the per-position correlation coefficients of sequence divergence versus inhibitor resistance, with the experimentally determined RDP for each inhibitor. The cut-off values obtained using the ROC analysis were 0.1700 for FPV and 0.1366 for SQV.

### Discussion

## Analysis of the identified positions

In this work, we found that in four heterogeneous protein families, a limited number of sequence positions are tightly linked to the studied phenotypes. This fact supports the notion that phenotypic adaptation can be achieved by a few sequence modifications. We reached this finding using an extremely simple strategy based on the correlation between sequence divergence and the quantitative phenotype differences, both obtained after pairwise comparisons. We developed a clustering heuristic which is based on the selection of a number of positions based on the *ρ*-value obtained when they are jointly analyzed (Figure 1B). This aided us in defining a set of residues associated with the explored phenotypes. As we show in the *Results* section, most of the positions identified by PhISCO were previously described as associated with function. For three of the studied cases, positions in direct interaction with ligands appear at the top of the scoring list. This is the case for Adk, Br and HIV-PR. Noteworthy, PhISCO also detects positions that are not in close proximity to the functional or catalytic sites. A rather indirect effect captured by our method can be mentioned for myoglobin, where there is a correlation between the net surface charge and its muscle concentration. Position 101 of this protein does not involve charged residues in our dataset, however it can still be associated with myoglobin concentration modulation. Interestingly, in close proximity to position 101 is the charged position 152. This suggests an interaction between both residues, where specific changes at position 101 may be required in order to accommodate a charged residue at position 152 without destabilizing the protein. It should be noted that our methodology for finding sequence positions associated with phenotype variations does not take explicitly into consideration any epistatic effect between sites. However, positions linked to non-additivity effects may be captured. The simplicity of the heuristic chosen to find positions make PhISCO unable to differentiate between additive and non-additive positions. For example, in the case of HIV-PR the non-additive effect of the mutations at PhISCO-identified positions 89 and 90 (where mutations L to V and L to M occur, respectively) has been thoroughly studied in both a theoretical and experimental manner [34]. This may be of particular relevance for phenotype prediction. Moreover, figure S2B and S2E suggest that the association between divergence and the differences in the antiviral drug inhibition is not linear, giving room to further improvements in the phenotype prediction by including epistatic effects. A similar epistatic effect could be pointed out in the case of residue 175 of AdK which is identified by PhISCO. Couñago et al. showed that the equivalent position in the *Bacillus subtilis* AdK (residue 179) is involved in protein adaptation to high temperatures [15]. The mutation T179I has a minor but significant effect in stability in the wildtype background sequence, while it has a significant stabilizing effect when it occurs in association with the Q199R mutation, which has no significant effect by itself. Although the found positions have been previously documented to be involved in enzyme function modulation and that these positions are used with success to predict the phenotype, conclusions should be taken with care. Due to the phylogenetic structure of the organisms involved in the analysis, some neutral mutations may arise as a consequence of a fixation in the different lineages. The actual version of PhISCO is not taking into account this effect.

Since the positions associated with resistance in HIV-PR are extensively characterized, this case gave us the opportunity to test our method in terms of sensitivity and specificity. In the first place, it was observed that most of the experimentally known RDPs appear among those best scored by PhISCO. In addition, assessing the cut-off value provided by the ROC analysis, which represents an equal weight of specificity and sensitivity, it was possible to adequately classify the correlated positions as RDP, with an accuracy of 90 %. Interestingly, the clustering strategy by PhISCO, based on the sequential addition of best scoring positions, resulted in selection of 7 positions, all of them corresponding to RDP. This result suggests that our method to analyze joint correlation favors specificity over sensitivity.

## Comparison to the state-of-the art methodologies

Although there is a myriad of available methods that do phenotype predictions, our proposed algorithm is one of the few that can be used for a diverse set of quantitative phenotypes. We designed PhISCO as a simple and interpretable algorithm for the prediction of quantitative phenotypes. Compared to other existing methods, both PhISCO and Signisite [13] allow obtaining positions associated with the phenotype by studying the relationship between a set of aligned sequences and their corresponding phenotype, expressed as a continuous quantifiable variable. Signisite analyzes the residue composition at a given position based on the magnitude of its associated phenotype; briefly, it orders the sequences according to their phenotype and analyzes the abundance of each amino acid according to its position in the ranking. On the other hand, PhISCO identifies positions performing a correlation of its sequence divergence with its phenotype difference. The performance of both methods can be compared in the case of HIV-PR, where the relevant positions for modulating the phenotype are known. As we can see in Table S2, both methods perform similarly in terms of accuracy and specificity, with PhISCO’s sensitivity being remarkably higher when using the cutoff derived from the ROC analysis. Although both methods are simple and fast, PhISCO adds a new capability which is the ability to make predictions of the phenotype value for sequences with unknown phenotype values. We compared our method with some of the available state-of-art strategies. For OGT prediction, the method of Li et. al. [11] which is a machine learning algorithm based on full proteome information, attains a RMSE (root-mean squared error) value of 2.16 °C and R^2^ of 0.95. The multiple linear regression method reported by Sauer et. al. [12], which uses several genome-derived features, attains a RMSE of 4.93 °C and R^2^ of 0.767. Even though both methods perform better than PhISCO in OGT prediction (RMSE of 8.5 °C and R^2^ of 0.85), it should be remembered that the performance obtained by this method is attained by the solely use of a protein family (Figure S5). In the case of predictions of absorption wavelengths of bacterial rhodopsins, PhISCO performs with a MAE (mean-absolute error) of 17.3 nm and R^2^ of 0.72, which is in the same order as for the machine learning-based method published by Karasuyama et al. (MAE of 7.8 nm and R^2^ of 0.774 and MAE of 12.4 nm and R^2^ of 0.8594, for 2 different rhodopsin subgroups). Although performance is better in the case of the Karasuyama et al., it comes with a cost of expensive training of thousands of parameters of a machine learning algorithm. Also, this training is performed for a specific protein family as a difference with PhISCO which is easily tunable for different protein families and phenotypes (Figure S6). Moreover, when studying the inhibitory efficiency for molecules that bind the HIV-PR we found that Beerenwinkel et al. developed the Gene2Pheno [42] algorithm based on support vector machines and perform predictions with a MSE = 0.204 and R^2^ = 0.71 for SQV inhibitor. PhISCO yielded a MSE = 0.21 with a R^2^ = 0.73. Although the cited publication is 20 years old, we were not able to find a recent software that performs this prediction for a similar dataset. It should be remembered at this point that most of the algorithms perform position finding or phenotype classification. In this sense, PhISCO do both position finding and phenotype prediction. For the case of myoglobin, no prediction method of its muscular concentration is available as far as we know.

## Concluding remarks

Since PhISCO identify positions tightly linked to phenotypes and, using a linear model, predict unknown phenotypes for target sequences with reasonable accuracy, it suggests that predictions can be performed with no explicit epistatic considerations. PhISCO uses a minimal set of parameters for predictions (the two coefficients of linear regression) which reflect an acceptable accuracy level even when a very limited number of data in the training set is available. Also, predictions performed using the whole sequences were clearly worse, which reinforces the idea of phenotypes being mostly codified in small subset of residues. It is interesting that the identity matrix implemented is the simplest penalization scheme, however, more complex amino acid substitution matrices may lead to similar or better results, although this was not explored in this work.

Overall, PhISCO is a generalist algorithm that performs at a level comparable to machine-learning based methods without the need for extensive training. Also the method is really simple since predictions are obtained at a very low information cost with results comparable to the state-of-art methodologies which are, in most of the cases, far more complex. In sum, we consider PhISCO a successful first-generation interpretable algorithm for the prediction of quantitative phenotypes and the associated sequence positions.

### Materials and methods

## Sequence datasets with their associated-phenotypes

To study temperature adaptation in Adenylate kinases (Adk), we used a database of Optimal Growth Temperatures (OGTs) compiled by our group, consisting in Archaeal organisms with fully sequenced genomes, which is an extended version of the data contained in Aptekmann et al. [14]. To this end, OGT values were collected from published articles identified from PubMed searches using the organism’s name as a query. When no OGT was available, it was approximated as the mean of the reported minimal and maximal growth temperatures of the organism. Available AdK protein sequences from the organisms in the definitive database were collected using the blastp search tool [38]. Multiple sequence alignments (MSA) were performed using the online server MAFFT with default settings. Columns with more than 10% of gaps were eliminated. As another study case, we studied a set of sequences of microbial rhodopsins with their light-absorption wavelengths obtained from *Karasuma et al.* [8]. The alignment and trimming strategies were the same as in the case of AdK. In a third example, we worked with a set of extant mammalian myoglobin sequences whose alignment and maximal muscle concentration were obtained from Mirceta et al. [29]. Finally, we studied sequences of HIV protease from clinical isolates and their associated change in the protease inhibitor drug resistance. This information was derived from the Stanford University HIV Drug Resistance Database (HIVdb), as published in Rhee et al. [40] (accessed in February 2022). Due to the high sequence conservation in HIV protease datasets, alignment and gap placement were unnecessary. Phenotypes refer to the level of protease-inhibition resistance, expressed as fold changes of inhibition (the ratio of concentration necessary to inhibit replication of a given strain compared with the wild type). The list of drug resistance positions (DRP) was obtained combining data from the Protease Inhibitors Resistance Notes of HIVdb available at https://hivdb.stanford.edu/dr-summary/resistance-notes/PI and the 2019 edition of the IAS–USA drug resistance mutations list [41].

Cluster analysis for all the studied cases were performed using CLANS [39], a clusterization algorithm based on *E*-values from pairwise-sequence comparisons, using a Network based method with global average.

## Phenotype predictions

With the aim to evaluate the performance of PhISCO’s phenotype prediction, the input sequences were assigned to two groups: a training set and a validation set. For a given position or set of positions, the relationship between sequence divergence for a pair i and j from the training set and its observed ΔPhenotype was modeled according the following linear equation:

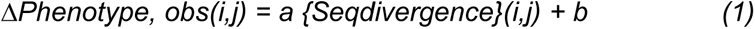

Each k sequence from the validation set was compared to the training sequences (j) yielding as many pairs of sequence divergence values as the number of sequences in the training set. From these divergences, the values of the predicted ΔPhenotype were calculated using the fitted coefficients of Equation 1, according to:

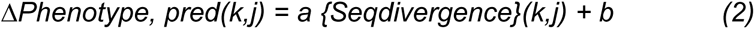

To do the prediction, the target phenotype was chosen as the one that minimizes the squared-sum of the difference between the predicted ΔPhenotype from the linear model (ΔPhenotype,pred(k,j)) and the difference between the unknown (Phenotype,pred(k)) and each known phenotype from the training set (Phenotype,obs(j)):

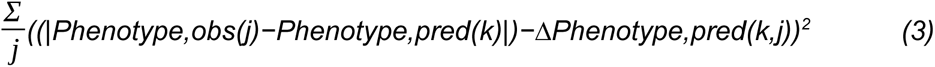

## Receiver operating characteristic (ROC) equations

*For ROC analysis in the HIV-PR case, the number of correctly predicted RDPs and non RDPs are indicated as true positives (TP) and true negatives (TN), whereas the number of incorrectly predicted RDPs and non RDPs are indicated as false negatives (FN) and false positives (FP), respectively. From these values, we calculated Sensitivity and Specificity, as defined below:*

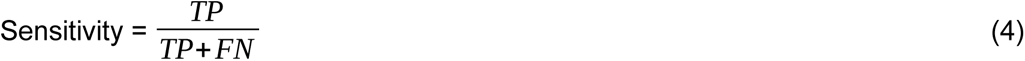

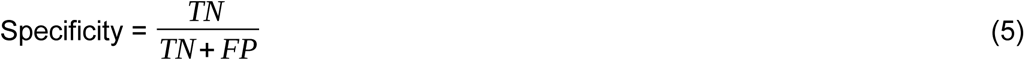

*The receiver operator characteristic (ROC) curve was derived from these measures calculated at different cutoff values (for review see [43]) and from that we obtained the area under the ROC curve (AUC). An “optimal” cutoff point was calculated from the maximal Youden’s index (J), defined as:*

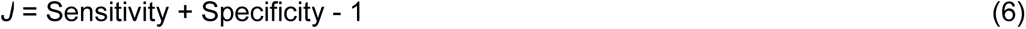

*Finally, the accuracy and the Matthews correlation coefficient were calculated as follows:*

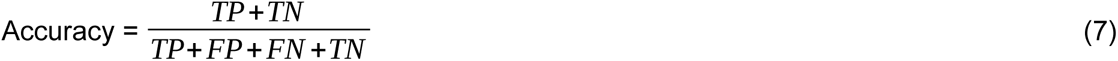

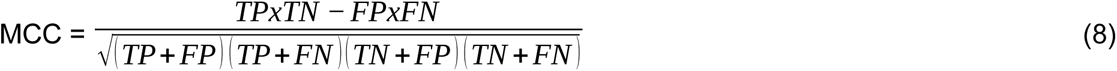

## Data analysis and scripts deposition

The calculations scripts were developed using standard python libraries and an implementation using Jupyter Notebooks. Both these scripts and the MSAs are deposited in the following repository: https://doi.org/10.5281/zenodo.7761213.

## Accession numbers

PDB ID **1AKE**, PDB ID **1C3W**, PDB ID **1ABS,** PDB ID **3OXC,** PDB ID **3EKV.**

## **Acknowledgements**.

This work was funded by the *CONICET* (PIP 2017) and *Agencia Nacional de Promoción de la Investigación, el Desarrollo Tecnológico y la Innovación* (PICT 2016 0014). E.A.R. thanks M.D. Ph.D. Lázaro Efraín Gerschenson for always trying to do the best he can with what he has.

