## Supplementary Information for "PhISCO: a simple method to infer phenotypes from protein sequences"

^f^ Universidad de Buenos Aires. Consejo Nacional de Investigaciones Científicas y Técnicas. Instituto de Química Biológica de la Facultad de Ciencias Exactas y Naturales (IQUIBICEN). Facultad de Ciencias Exactas y Naturales. Laboratorio de Fisiología de Proteínas. Buenos Aires, Argentina

^g^ Consejo Nacional de Investigaciones Científicas y Técnicas, Instituto de Química y Fisicoquímica Biológicas, Buenos Aires, Argentina

^h^ Departamento de Ciencia y Tecnología, Universidad Nacional de Quilmes, Argentina

^I^ Department of Biochemistry and Microbiology, Rutgers University, New Brunswick, NJ 08873, USA

^J^ Institute of Marine and Coastal Sciences, Rutgers University, New Brunswick, NJ 08901, USA

*****Both authors contributed equally to this work

**To whom correspondence should be addressed: .

Running title: Inferring phenotypes from sequences.

#### Table S1. Top 20 scoring positions for resistance of HIV-PR to PI

| **FPV** | | | **SQV** | | |
| --- | --- | --- | --- | --- | --- |
| **Position** | **Score** | **RDP** | **Position** | **Score** | **RDP** |
| 84 | 0.502 | + | 84 | 0.505 | + |
| 54 | 0.497 | + | 54 | 0.446 | + |
| 46 | 0.458 | + | 71 | 0.428 | + |
| 10 | 0.426 | + | 90 | 0.420 | + |
| 33 | 0.417 | + | 10 | 0.384 | + |
| 32 | 0.391 | + | 48 | 0.319 | + |
| 47 | 0.356 | + | 73 | 0.294 | + |
| 71 | 0.316 | - | 33 | 0.277 | - |
| 90 | 0.303 | + | 46 | 0.262 | + |
| 73 | 0.290 | + | 20 | 0.255 | + |
| 11 | 0.279 | + | 53 | 0.236 | + |
| 82 | 0.273 | + | 82 | 0.211 | + |
| 20 | 0.233 | + | 74 | 0.171 | - |
| 43 | 0.210 | - | 32 | 0.156 | - |
| 76 | 0.208 | + | 11 | 0.150 | - |
| 53 | 0.190 | - | 34 | 0.149 | - |
| 58 | 0.176 | - | 24 | 0.137 | + |
| 34 | 0.175 | - | 66 | 0.135 | - |
| 24 | 0.170 | + | 43 | 0.131 | - |
| 91 | 0.165 | - | 36 | 0.126 | - |

**Table S2.** Comparison between PhISCO and SigniSite performance^1^.

|  | *SigniSite* | *PhISCO* | |
| --- | --- | --- | --- |
|  |  | *Heuristic^2^* | *ROC^3^* |
| *Accuracy* | *0.90* | *0.89* | *0.90* |
| *Sensitivity* | *0.61* | *0.45* | *0.84* |
| *Specificity* | *0.98* | *1.00* | *0.92* |
| *MCC* | *0.68* | *0.63* | *0.72* |

*^1^Statistics for both analysed HIV-PR inhibitors (FPV and SQV) are averaged.*

*^2^PhISCO cut off value derived from the heuristic to obtain the set of positions that give the highest correlation in Figure S2.*

*^3^PhISCO cut off derived from the ROC analysis in Figure 5.*

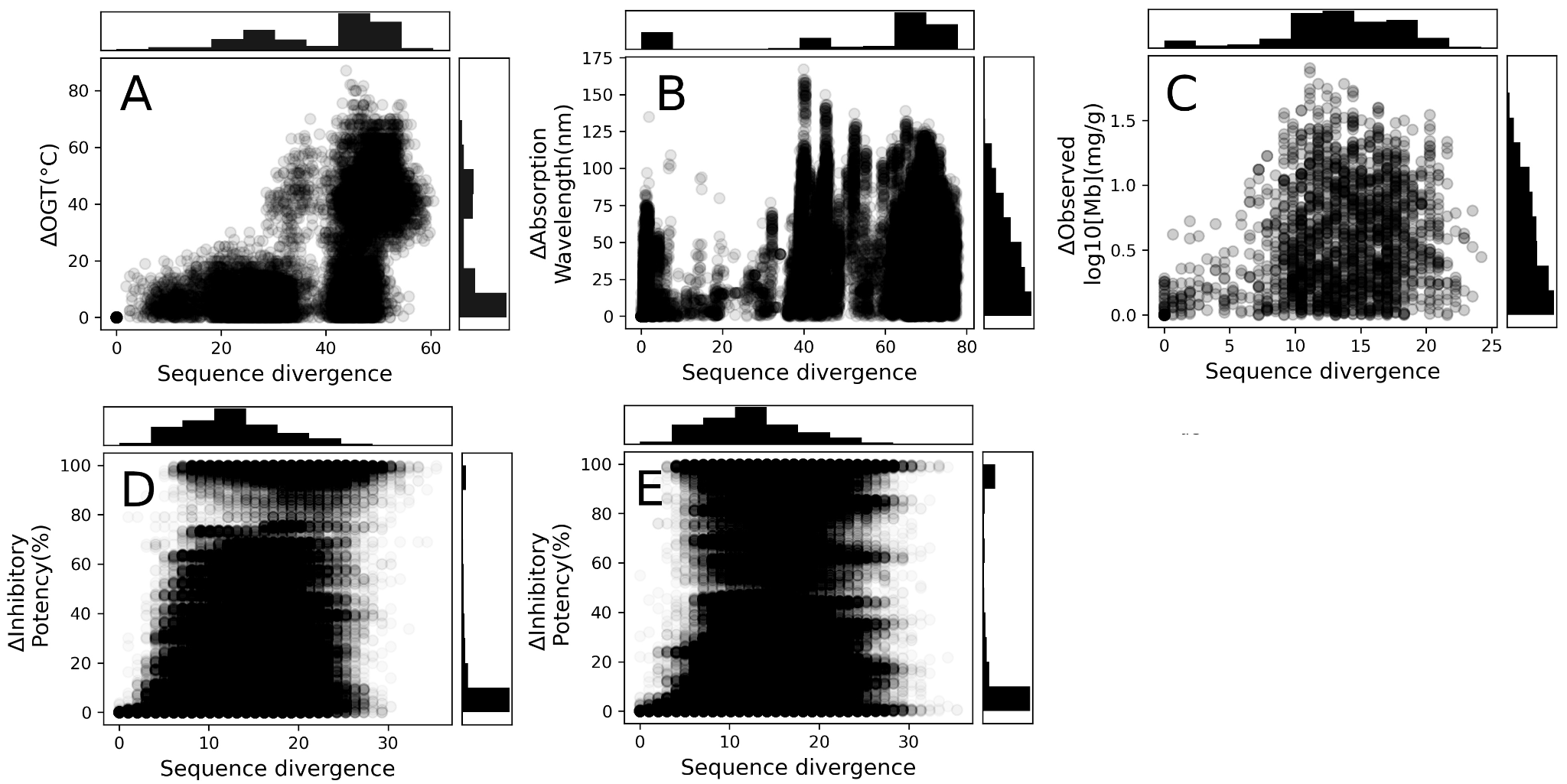

#### Figure S1. Whole sequence divergence and *ΔPhenotypes.* The upper panel is the histogram of the sequence divergence while the right panel is the histogram of the *ΔPhenotype.* A. The case of AdK and the organisms’ optimal growth temperature. B. The case of Br and the maximal absorption wavelength. C. The case of mammalian divers and their myoglobin muscle concentration. D and E. The case of HIV protease and the inhibitor potency of FPV and SQV, respectively.

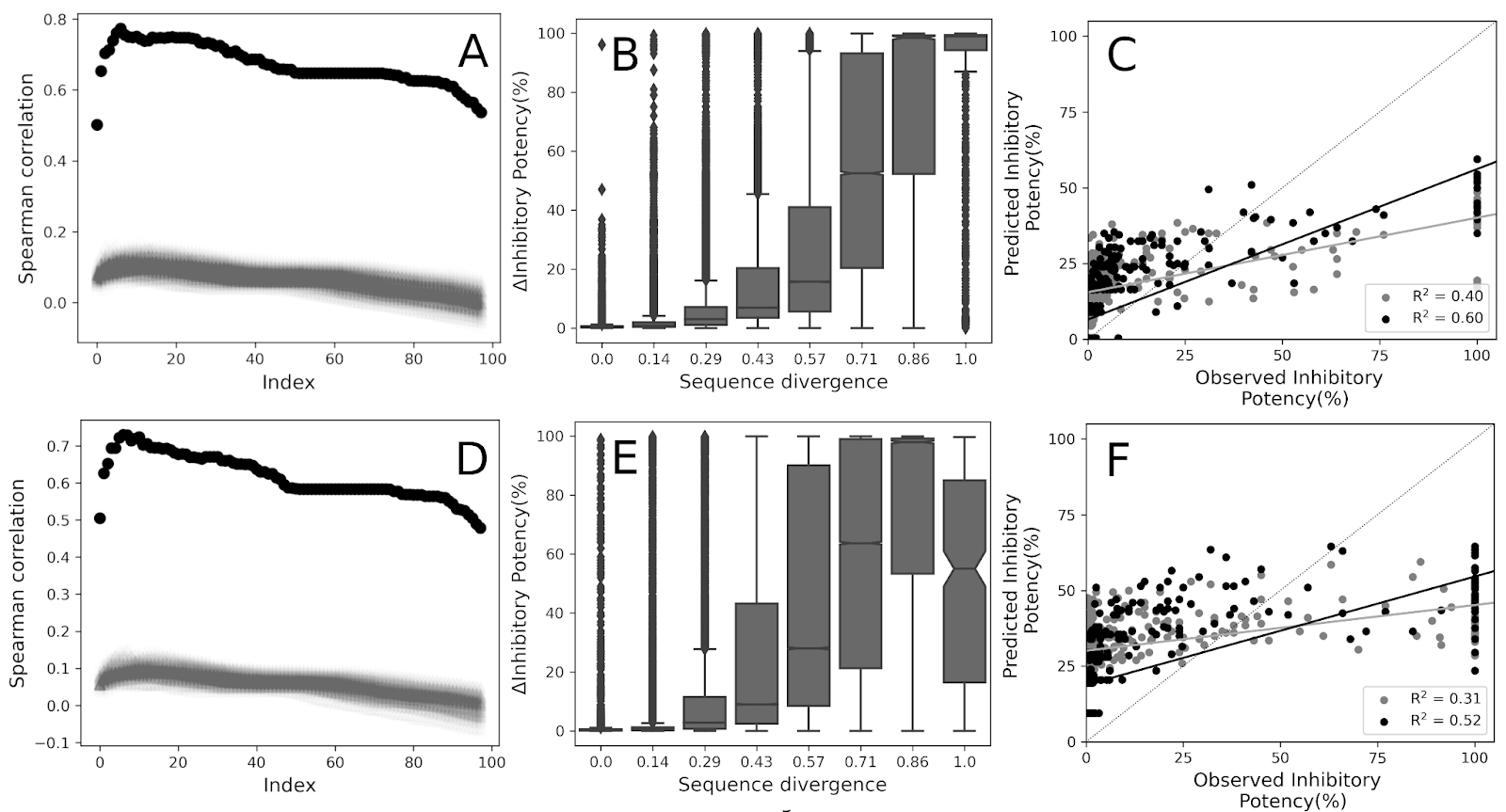

**Figure S2.** Inhibitory potency of Fosamprenavir (FPV) and Saquinavir (SQV) in clinical isolates of HIV-protease (Hp). A and D. Correlation coefficients of the Hp individual positions in a descendent order (black) and of the accumulated scores in the same order (red) for FPV and SQV, respectively. B and E. *E. Distribution of the phenotype differences at different sequence divergence values. (R2 = 0.66  and 0.59 from a linear fit of inhibitory potency of FPV and SQV vs sequence divergence)*. C and F. Prediction of logarithm of the inhibitory potency for FPV and SQV, respectively, from sequences in one validation set chosen to have a representative Pearson correlation coefficient associated.

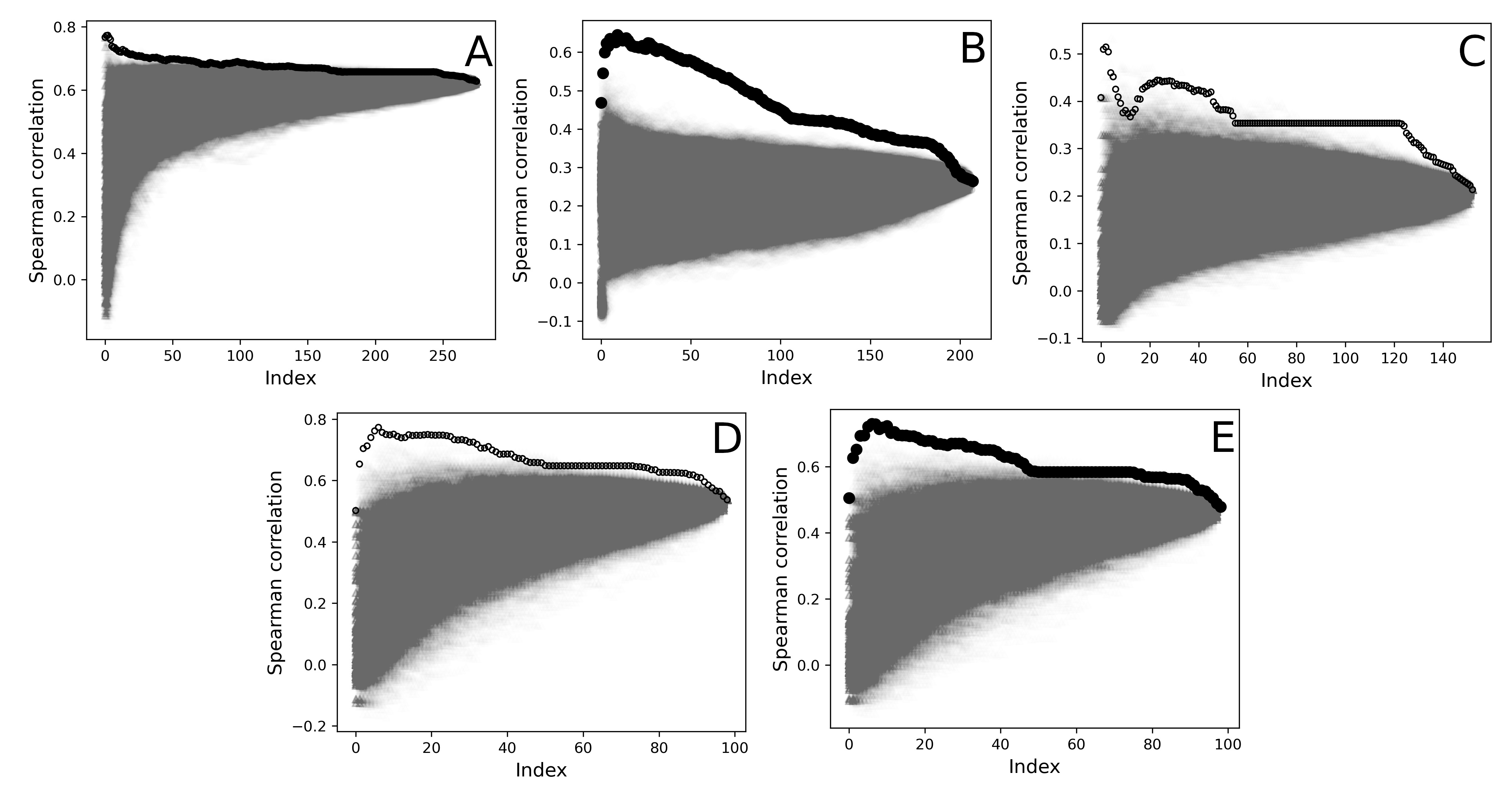
*Figure S3. Control calculations for PhISCO’s joint correlation analysis. Correlation coefficients by addition of positions ranked in a descendant order (black filled circles) or after the addition of randomly selected positions (5000 rounds, gray points).* ***A****. The case of AdK.* ***B.*** *The case of microbial rhodopsins.* ***C.*** *The case of muscle myoglobin.* ***D.*** *The case of inhibition of HIV-PR with Fosempavir.* ***E.*** *The case of inhibition of HIV-PR with Saquinavir****.***

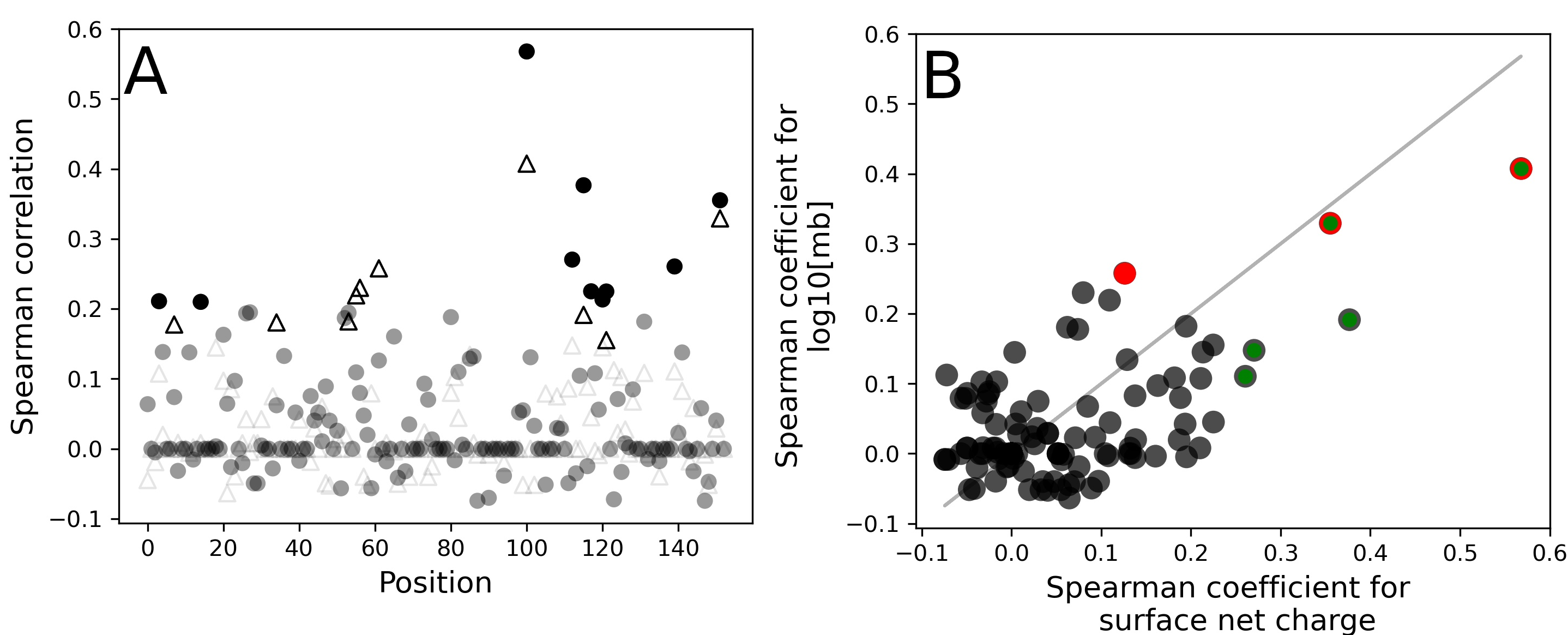

*Figure S4. Positions involved in different traits associated with myoglobin.* ***A.*** *Solid circles are the rho values for the correlation between sequence divergence and muscular myoglobin concentration, while empty triangles are the rho values for the correlation between sequence divergence and myoglobin net surface charge.* ***B.*** *Spearman coefficients for the correlation between sequence divergence and muscular myoglobin concentration (gray circles) against the rho values for the correlation between sequence divergence and myoglobin net surface charge (black circles). The red symbols are the positions found by PhISCO in the joint correlation analysis to predict myoglobin concentration while green circles are the ones found by PhISCO to predict the net surface charge.*

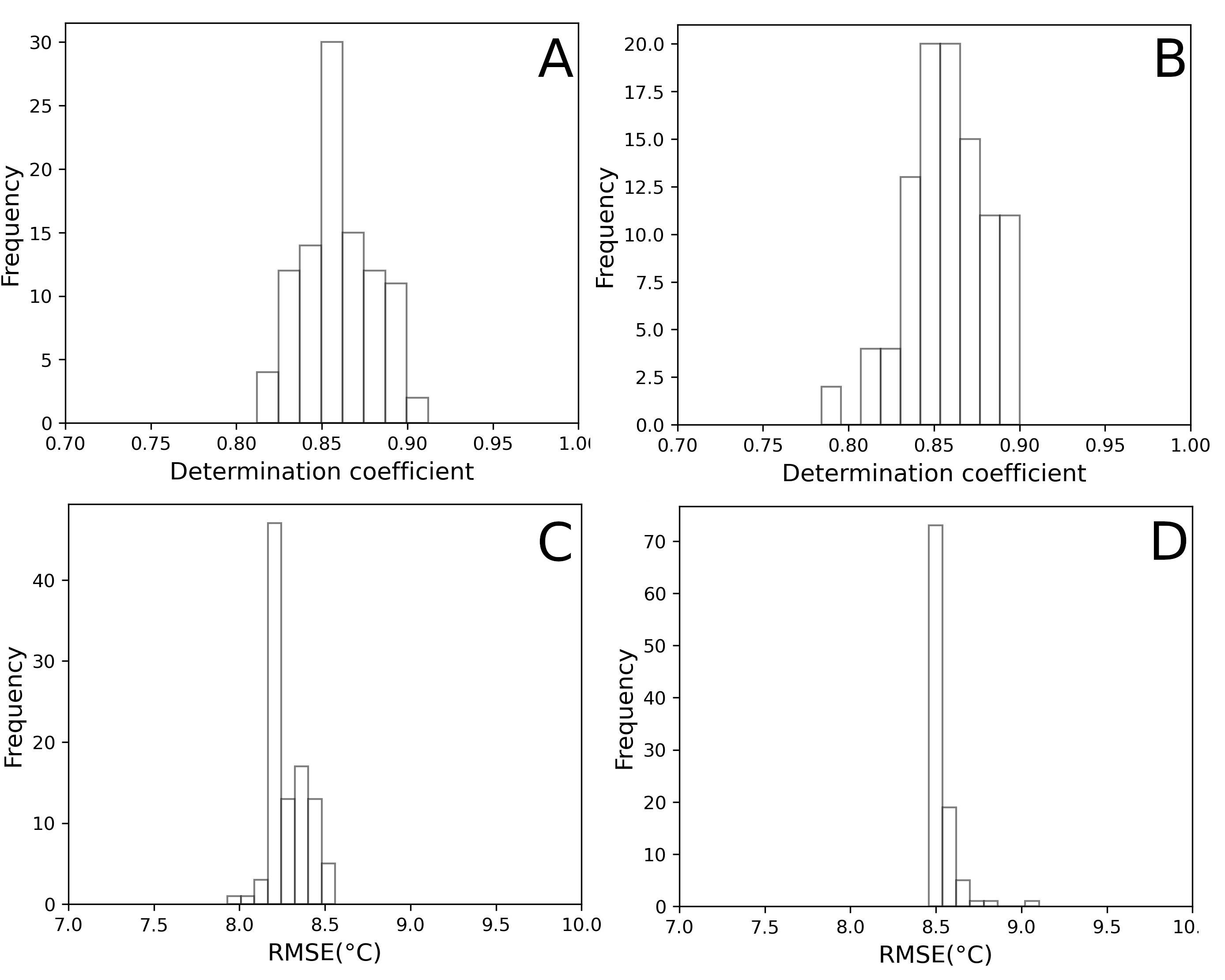

*Figure S5. Performance of PhISCO predictions in 100 random selected training and validation datasets for the ADK case.* ***A and B****. Distribution of R2 for training and validation sets, respectively.* ***C and D****. Distribution of root-mean squared errors (RMSE) for each prediction round (training and validation sets, respectively).*

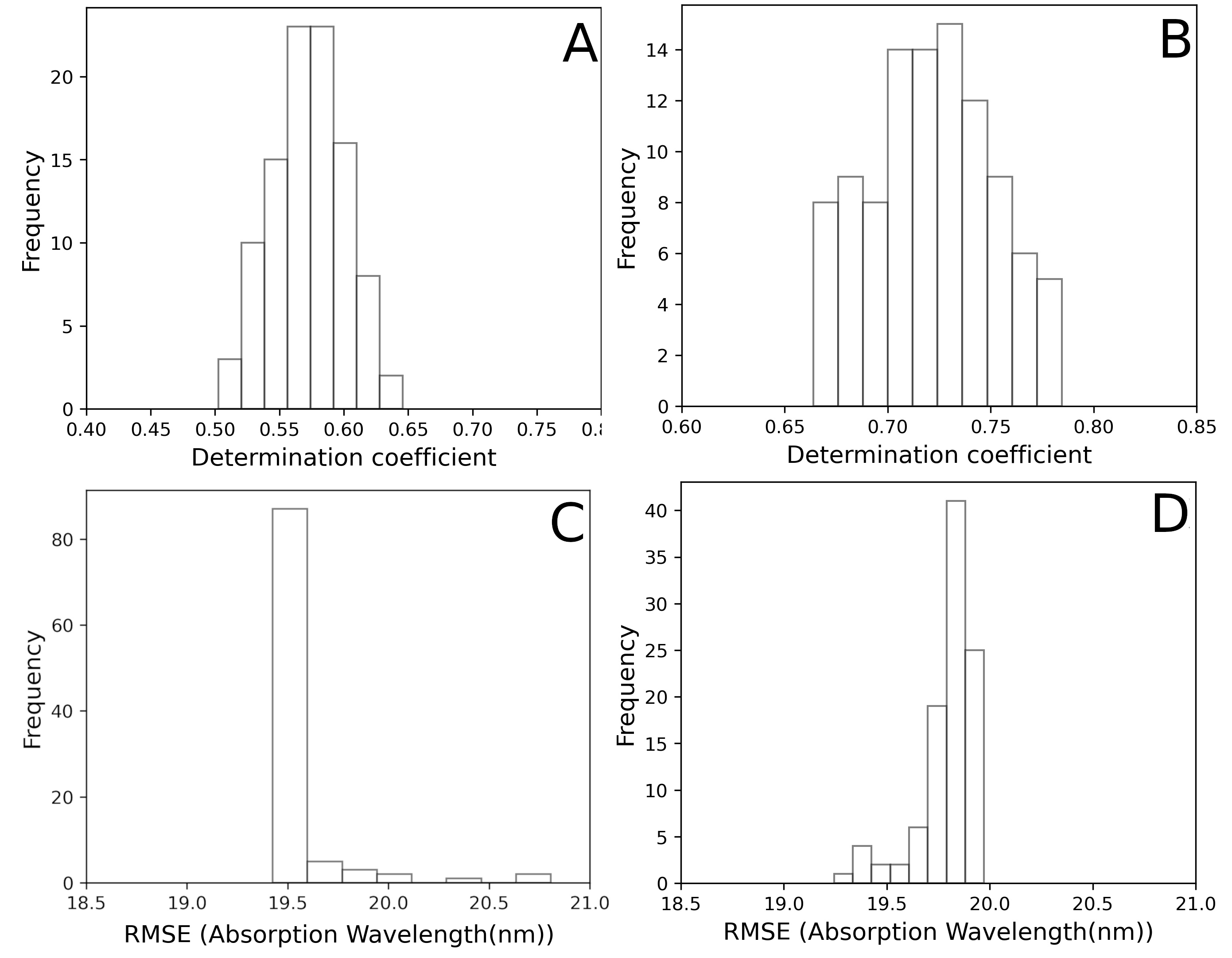

*Figure S6. Performance of PhISCO predictions in 100 random selected training and validation datasets in the microbial rhodopsins case.* ***A and B****. Distribution of R2 for training and validation sets, respectively.* ***C and D****. Distribution of root-mean squared errors (RMSE) for each prediction round (training and validation sets, respectively).*

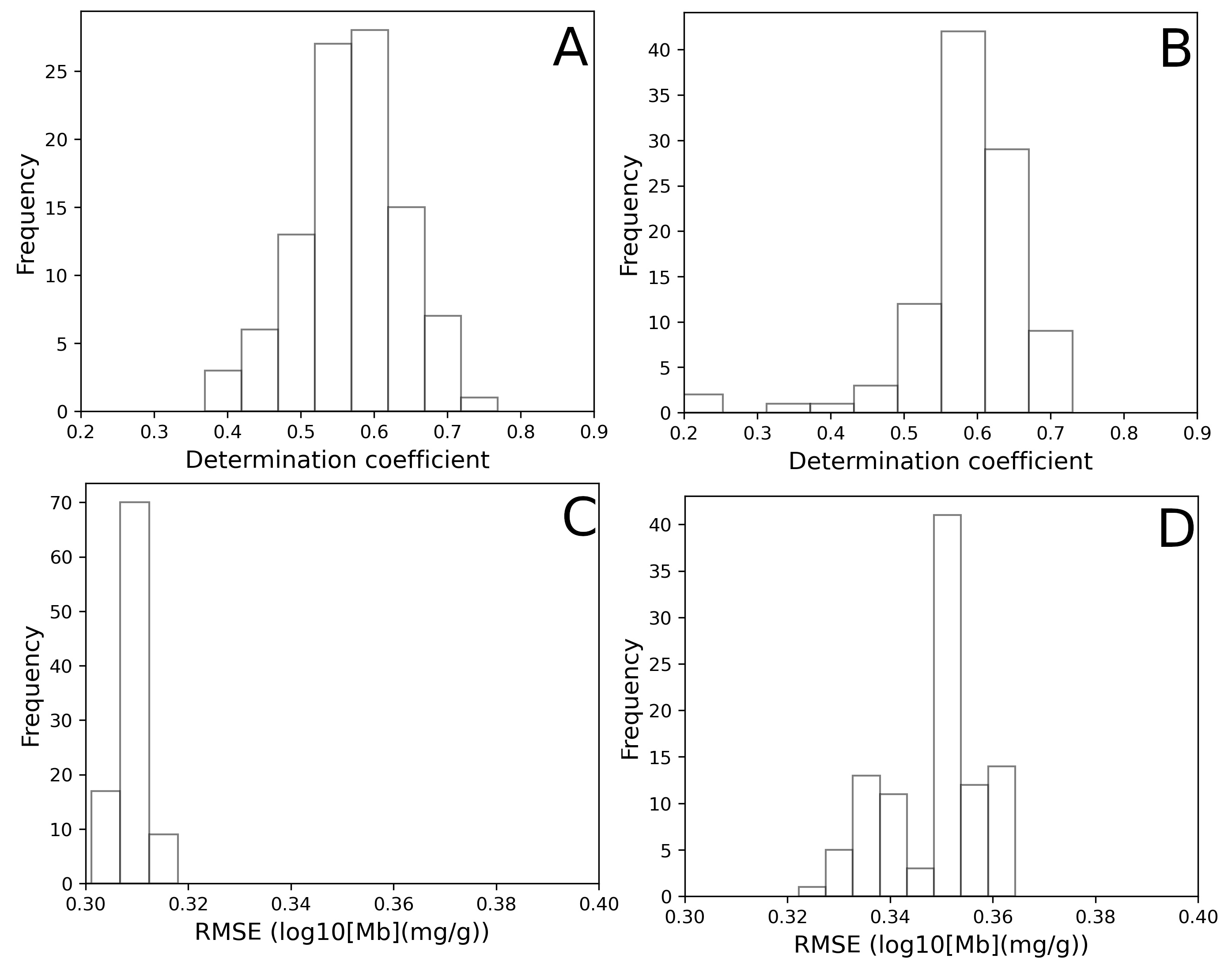

*Figure S7. Performance of PhISCO predictions in the 100 random selected training and validation datasets in the myoglobin case.* ***A and B****. Distribution of R2 for training and validation sets, respectively.* ***C and D****. Distribution of root-mean squared errors (RMSE) for each prediction round (training and validation sets, respectively).*

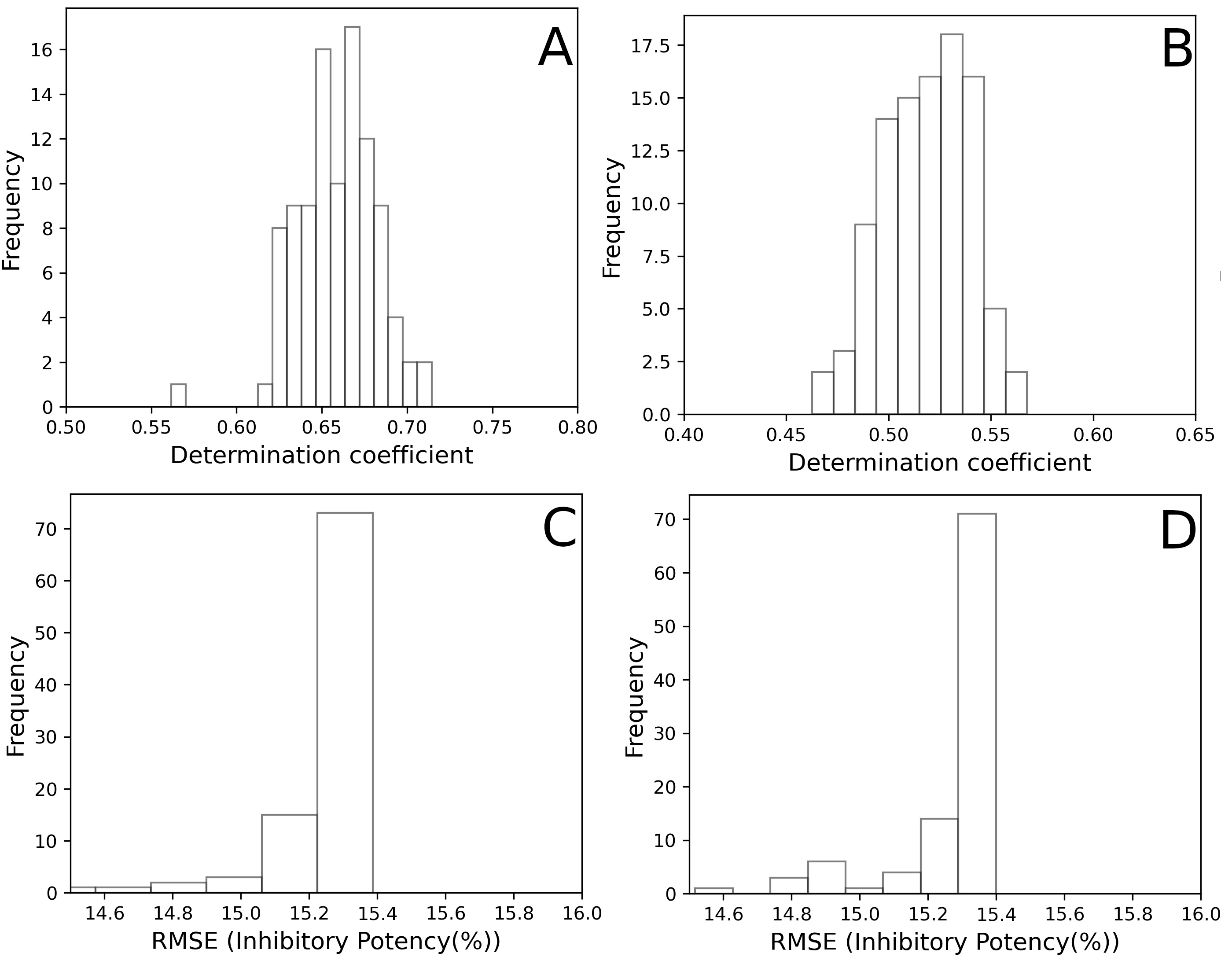

*Figure S8. Performance of PhISCO predictions in the 100 random selected training and validation datasets for HIV-PR inhibition with FPV.* ***A and B****. Distribution of R2 for training and validation sets, respectively.* ***C and D****. Distribution of root-mean squared errors (RMSE) for each prediction round (training and validation sets, respectively).*

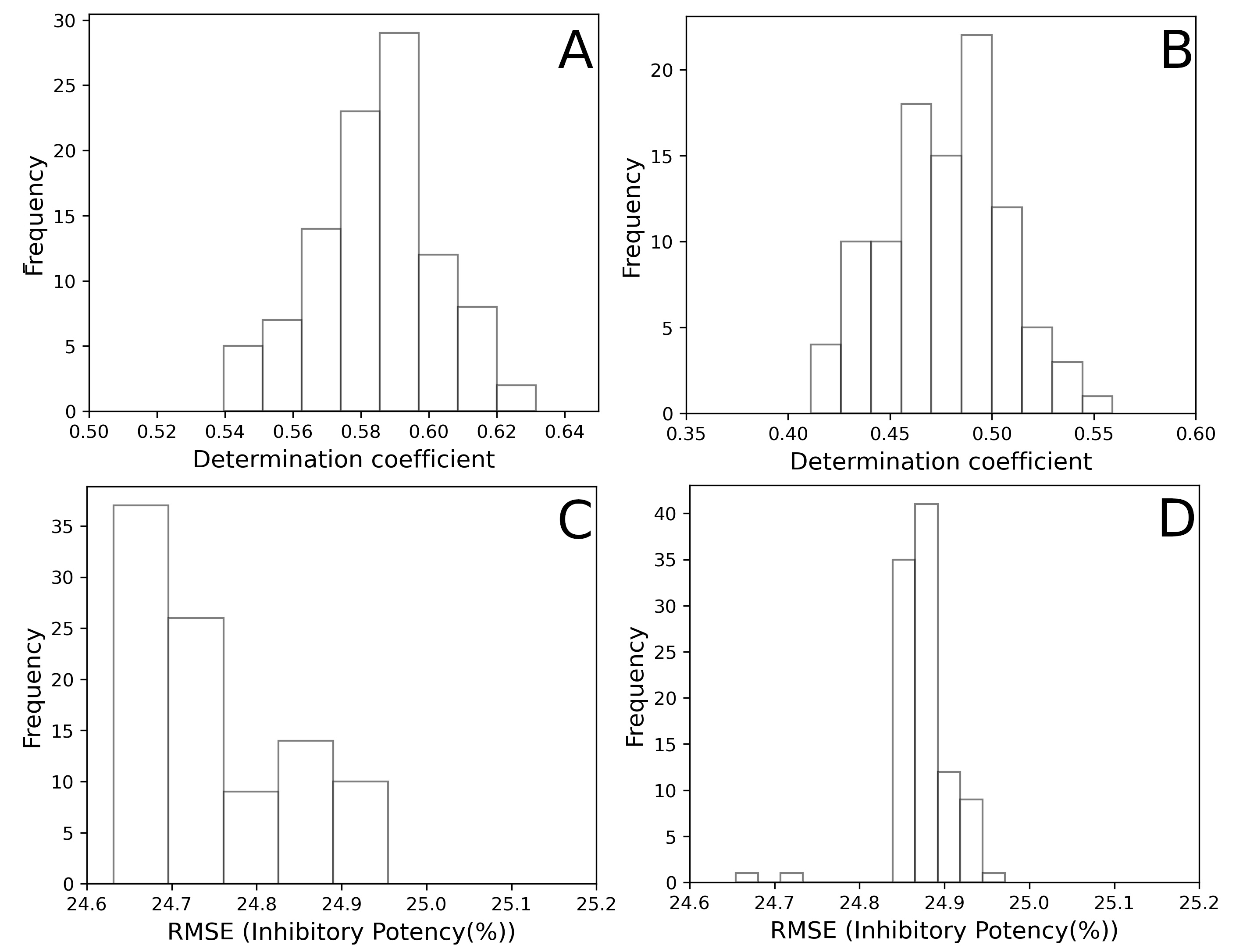

*Figure S9. Performance of PhISCO predictions in the 100 random selected training and validation datasets for HIV-PR inhibition with SQV.****A and B****. Distribution of R2 for training and validation sets, respectively.* ***C and D****. Distribution of root-mean squared errors (RMSE) for each prediction round (training and validation sets, respectively).*

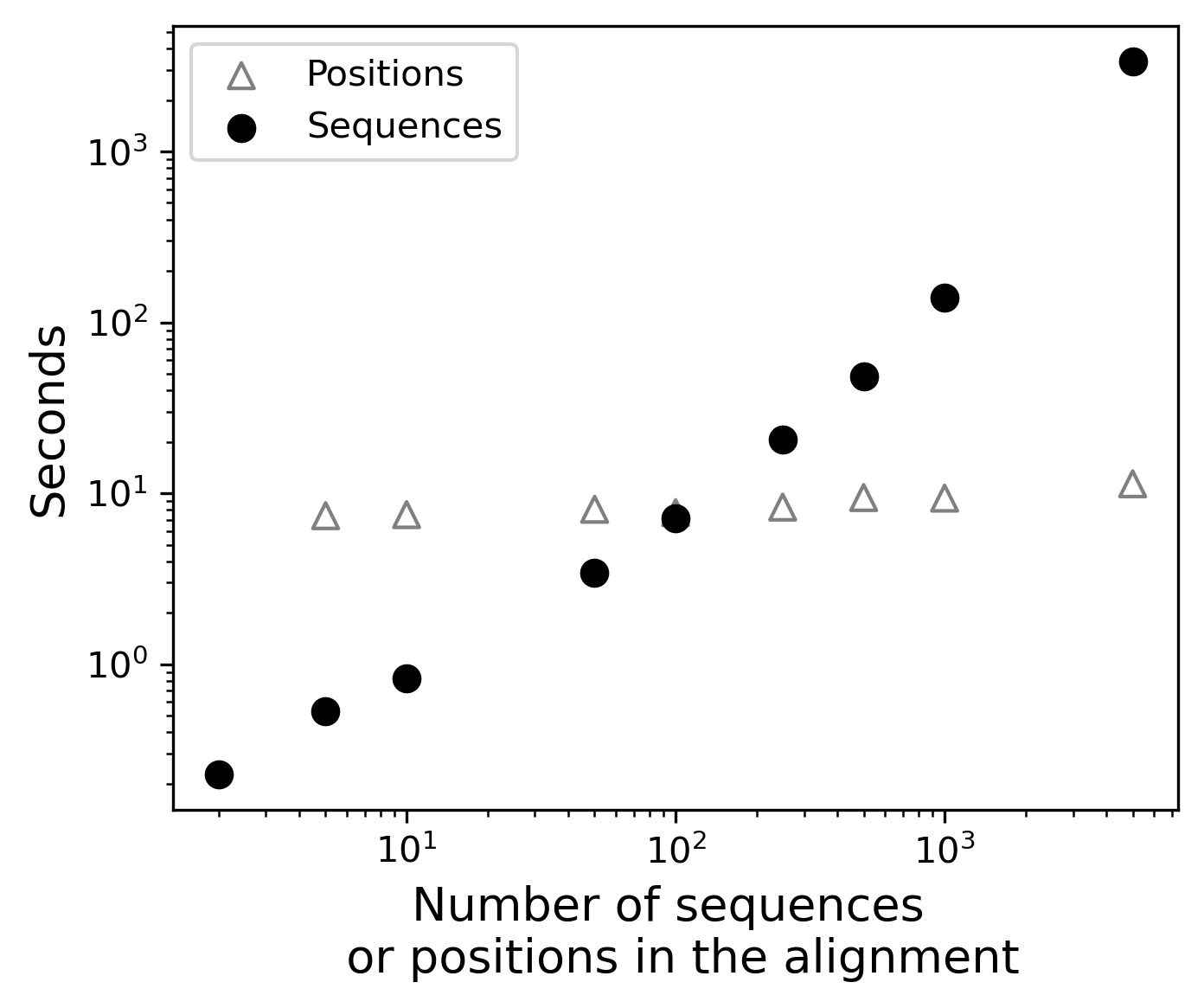

*Figure S10. Performance of PhISCO in an alignment with 100 sequences and different number of positions (open triangles) and with 100 of positions and different number of sequences (black circles).*
